# Few-shot regulatory sequence design with a genomic language model

**DOI:** 10.64898/2026.10.28.762639

**Authors:** Yukun A. Hao, Matthew Liu, Divya Nori, Garyk Brixi, Thomas Lau, Charu Ramakrishnan, Daniel Chang, Ryan Park, William E. Allen, Brian L. Hie

## Abstract

A central challenge in generative biology is translating knowledge learned from natural sequences and experiments into designs with desired functions. In-context learning (ICL) is an emergent capability of large language models that allows them to adapt to new tasks from a few examples without additional training, enabling these models to become general-purpose learners. Here, we show that the genome language model, Evo 2, exhibits generative ICL in biological sequence design. Prompting with just tens of examples conditions Evo 2 to generate novel and diverse regulatory DNA sequences without any additional training. Mechanistic analysis suggests that genomic ICL may originate from induction, similar to phenomena observed in natural language. Reinforcement learning leveraging sequence-to-function models as experiment-derived rewards further strengthens ICL. We first validate this approach by generating enhancers that drive cell-type-specific gene expression in three human cell lines. Extending this approach *in vivo*, enhancers designed using human regulatory element data selectively label layer 5 extratelencephalic neurons in the mouse cortex with high specificity and stronger activity than existing enhancers. Our study provides evidence of emergent properties in genomic sequence models and introduces a promising approach for improving functional conditioning in generative biology.

## Introduction

Composing artificial systems with desired functions is a central goal of synthetic biology. Generative genomic language models, such as Evo 1 and Evo 2, offer a unified foundation for designing DNA, RNA, and protein sequences by learning from the diversity of natural genomes (Brixi et al., 2026b; Nguyen et al., 2024). However, leveraging these models to design functional sequences still requires steering their generated outputs toward a specified set of functions. Prompting Evo with native genomic context can guide the generation of functional *de novo* genes in prokaryotic genomes (Merchant et al., 2026), while task-specific fine-tuning on thousands of examples can produce specialized sequence designers (King et al., 2026; Lal et al., 2024; Nguyen et al., 2024). However, natural genomic contexts or sufficiently large labeled datasets do not exist for every design objective. These limitations are especially restrictive for applications such as cell-type-specific enhancer design *in vivo*, where experimentally confirmed sequences are often scarce.

In-context learning (ICL), an emergent capability first demonstrated by GPT-3, established large language models as few-shot learners that can adapt to new tasks by using examples provided in a prompt, without updating their parameters (Brown et al., 2020). This could enable a new paradigm in generative biology in which a small set of functionally related sequences could itself specify a design objective. Recent work has documented some evidence of ICL in Evo 2 on out-of-distribution symbolic tasks (Breslow et al., 2025). However, whether this capability extends from symbolic pattern completion to the generation of sequences with defined biological function has not been tested.

Here, we show that Evo 2 can learn biological sequence design tasks from in-context examples, using eukaryotic regulatory sequences as a test bed. Without task-specific parameter updates, a single pretrained model generates novel sequences across five tasks including yeast promoters, human 5*^′^* untranslated regions (5*^′^* UTR), internal ribosome entry sites (IRES), human promoters, and human enhancers. Across diverse functional classes of regulatory elements, predicted design quality improves with more examples, indicating better task specification with more in-context demonstrations. By ablating specific attention heads in Evo 2, we observed circuits driving induction, providing a potential mechanism of genomic ICL.

Sequence-to-function models, neural networks trained to map DNA sequences to experimental measurements such as regulatory activity (Agarwal et al., 2025; Barbadilla-Martínez et al., 2026; Gosai et al., 2024; Pampari et al., 2025) or epigenomic features (Avsec et al., 2021, 2026; Linder et al., 2025) learn specialized knowledge of genomic structure and function (Cheng et al., 2025; Nagai et al., 2026), which can further augment generative models. We show that lightweight reinforcement learning that repurposes sequence-to-function as virtual rewards improves the predicted quality of sequences generated by genomic ICL. These improvements can transfer across regulatory classes and cellular contexts; for example, models trained for promoter design perform better on enhancer design, while those trained for cell-line-specific enhancer design generalize to a held-out cell line. Reporter assays in three human cell lines validate both the activity and specificity of Evo 2-generated enhancers and show that reinforcement learning can improve design quality.

We extend this approach by fine-tuning AlphaGenome (Avsec et al., 2026) on human brain chromatin accessibility data across 107 cell types (Li et al., 2023) and use its predictions to improve cell-type-specific ICL in Evo 2 through reinforcement learning. Two enhancers designed by this post-trained Evo 2 successfully label layer 5 extratelencephalic neurons in the mouse brain, demonstrating that this framework can yield functional tools for *in vivo* applications. Our findings establish genomic ICL as a practical approach to functional sequence design and show how reinforcement learning can strengthen this capability, providing a foundation for applying generative genomic models across a broad range of biological design tasks.

## Results

### Evo 2 learns biological sequence design from in-context examples

In-context learning enables pretrained natural language models to perform novel tasks specified by examples without updating their parameters, with more in-context examples typically leading to better task performance (Brown et al., 2020). We hypothesized that Evo 2, a generative genomic language model with a context window of up to one million base pairs, could similarly learn to generate novel biological sequences from prompted examples without further training. To test this, we concatenated DNA sequences with shared properties to prompt Evo 2 to generate additional sequences (**Figure 1A**).

**Fig. 1.**
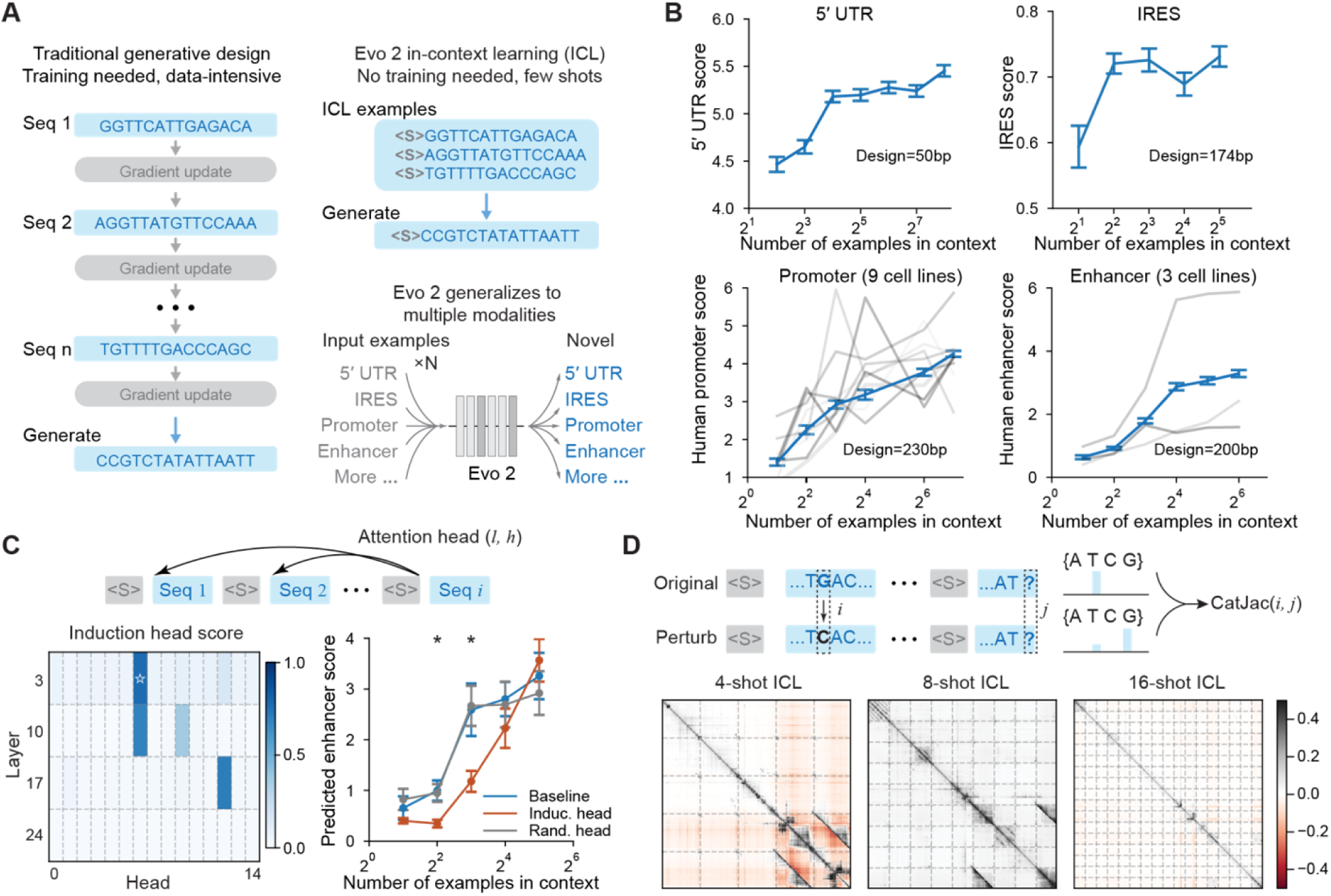
*|* **Evo 2 learns to design regulatory sequences from in-context examples. (A)** In-context learning (ICL) enables data-efficient sequence design across diverse regulatory tasks. In contrast to other methods, ICL does not require task-specific model updates and can leverage as few as tens of example demonstrations. **(B)** Predicted activity of sequences generated by Evo 2 through in-context learning for four tasks: human 5*^′^* UTR design, human IRES design, human promoter design, and human enhancer design. Error bars indicate the standard error of the mean (SEM; n = 128 generated sequences per condition). For human promoters, gray lines represent the nine cell lines, ordered from light to dark: AGS, HAP1, HCT116, HEK293, HepG2, K562, LNCaP, MCF7, and U2OS. For human enhancers, gray lines represent the three cell lines, ordered from light to dark: HepG2, K562, and SK-N-SH. **(C)** Identification and ablation of an induction attention head in Evo 2 1B. Left: induction-head score. The inductionhead score for head *h* in layer *l* is computed as the average score attending from the last token of a separating sequence to leading tokens of previous examples. The star marks the putative induction head, layer 3 head 6, selected for ablation. Right: predicted enhancer score of newly generated sequences after ablation of the induction head, ablation of a random attention head in the same layer, layer 3 head 7, or no ablation. Error bars indicate SEM (n = 32 ICL prompts per condition). The *p*-value for two-sided paired *t* -test is 0.023 (4-shot), 0.040(8-shot) after the Benjamini-Hochberg procedure. **(D)** Top: Dependency between generated sequences and prompted example sequences, measured using the categorical Jacobian (CatJac). Bottom: CatJacs are shown for 4-shot, 8-shot, and 16-shot in-context learning for human enhancer design.

As an initial experiment, we tasked Evo 2 with generating yeast promoter sequences. We constructed prompts containing 1, 2, 4, 8, 16, 32, or 64 highly active synthetic promoters that are 80 bp each and flanked by a fixed separator sequence. For each prompt, we generated 128 sequences and filtered out those with low complexity repeats, abnormal GC content, or high similarity to the prompt sequences and their reverse complements. We evaluated the remaining sequences using a neural network trained to predict yeast promoter activity (Lal et al., 2024). With as few as four examples, Evo 2 generated novel sequences with predicted activities comparable to those of highly active positive controls (**Figure S1A).** The predicted activity of generated promoters increased as Evo 2 was prompted with more examples before plateauing at *≥*8 in-context sequences. These results suggest that a small set of DNA examples can condition Evo 2 to generate novel sequences with similar predicted activity.

We next evaluated genomic ICL on human 5*^′^* untranslated regions (UTRs) and internal ribosome entry sites (IRESs). We applied the same prompt structure and filtering strategy to evaluate generated sequences using task-specific predictive models (Zhang et al., 2025; Zhao et al., 2024). For 50-bp 5*^′^* UTRs, we found that 16 examples were sufficient to condition Evo 2 to generate active sequences, with more than 40% passing a predetermined activity threshold. For 174-nt IRESs, 4-shot example prompting achieved a pass rate exceeding 60% (**Figure 1B, S1B**).

We then tested whether Evo 2 can learn to generate human regulatory sequences active in a specified cell line from in-context examples. For proximal regulatory sequences, we prompted Evo 2 with highly active, 200– 230-bp human promoters from each of nine cell lines and evaluated generated sequences using a published activity predictor after applying a similar set of novelty and validity filters (Barbadilla-Martínez et al., 2026). On average, more examples in the prompt conditioned Evo 2 to generate sequences with higher predicted activity in the specified cell type. With 128 examples in context, 50% of Evo 2-generated sequences were predicted to have a log2-fold-change of 4 (**Figure 1B**). For distal regulatory elements (Gosai et al., 2024), we observed similar ICL scaling when generating 200-bp human enhancers using examples active in each of three target cell lines (**Figure 1B**). Prompting Evo 2 with natural enhancer examples showed higher yield than prompting Evo 2 with their upstream genomic context (**Figure S2**), indicating that genomic ICL can be an effective way to condition Evo 2 on sequence design tasks. Taken together, these results indicate that in-context examples can specify multiple regulatory design tasks to Evo 2 (**Figure 1A**). More in-context examples yielded sequences with higher desired quality, establishing ICL scaling in generative genomic models.

### Specialized attention heads contribute to ICL in Evo 2

To investigate the underlying mechanism for in-context learning in Evo 2, we carried out two sets of causal perturbations. First, we looked for the existence of “induction heads,” which are attention modules that use repeated patterns to predict the next token (Olsson et al., 2022). Using duplicated random sequences and a smaller Evo 2 1B model, we identified attention heads that recognize repeated sequence content across the prompt. A subset of the heads also attended to earlier example sequences from later separators, suggesting Evo 2 recognizes the ICL prompt structure (**Figure 1C**). In enhancer design, ablating a candidate induction head reduced the average predicted activity of generated sequences when compared to intact and samelayer non-induction-head controls (**Figure 1C**). We similarly identified and ablated candidate induction heads in the larger Evo 2 7B model, and observed the same drop in predicted activity relative to ablating randomly chosen attention heads (**Figure S3**). This is consistent with the possibility that some attention heads are essential for inductive generation, an important finding when identifying circuits for in-context learning in large language models (Olsson et al., 2022). Moreover, using sparse autoencoders, a common interpretability tool (Huben et al., 2024) to learn sparse feature representations for Evo 2, we identified correlative features whose strength increases with more separator sequences or example sequences in-context, suggesting progressive internal recognition of the ICL prompt structure (**Figure S4**).

Second, we examined how individual examples in the context influenced generated sequences at nucleotide resolution using the “categorical Jacobian” method (Zhang et al., 2024). We mutated individual nucleotides in the concatenated prompt and generated sequences to quantify changes in the model-inferred nucleotide probabilities at other positions. In low-shot enhancer design across the three cell lines examined, high-influence nucleotides were often located in only a single example sequence (**Figure 1D**). However, as more examples enter the context, high-influence nucleotides become more distributed across the prompt (**Figure 1D**). This shift suggests that Evo 2 increasingly synthesizes information from multiple examples to inform ICL-based generation. Taken together, this data provides evidence that Evo 2 can perform inductive generation by synthesizing features across the examples in its context window.

### Experiment-derived rewards incentivize Evo 2 to design better regulatory sequences

Combining knowledge from experimental measurements with generative biological models could further align biological sequence design with desired functions (Widatalla et al., 2026). In natural language modeling, human feedback in the form of labeled preferences, neural network-simulated human feedback, and verifiable rewards are used to incentivize models to improve generated outputs (Guo et al., 2025; Ouyang et al., 2022). We hypothesized that we could apply sequence-to-function models, neural networks trained to predict experimental measurements from sequences alone, as virtual reward models to incentivize Evo 2 to design better sequences more efficiently using in-context learning. In particular, the sequence-to-function models trained on massively parallel reporter assays (MPRAs) can model the regulatory activity of sequences in defined biological contexts with high accuracy (Barbadilla-Martínez et al., 2026; Gosai et al., 2024; Tewhey et al., 2016). We combined these predictions with consensus constraints, such as GC content and novelty, to post-train Evo 2 through lightweight reinforcement learning (RL) to improve the sequences generated by ICL (**Figure 2A**).

**Fig. 2.**
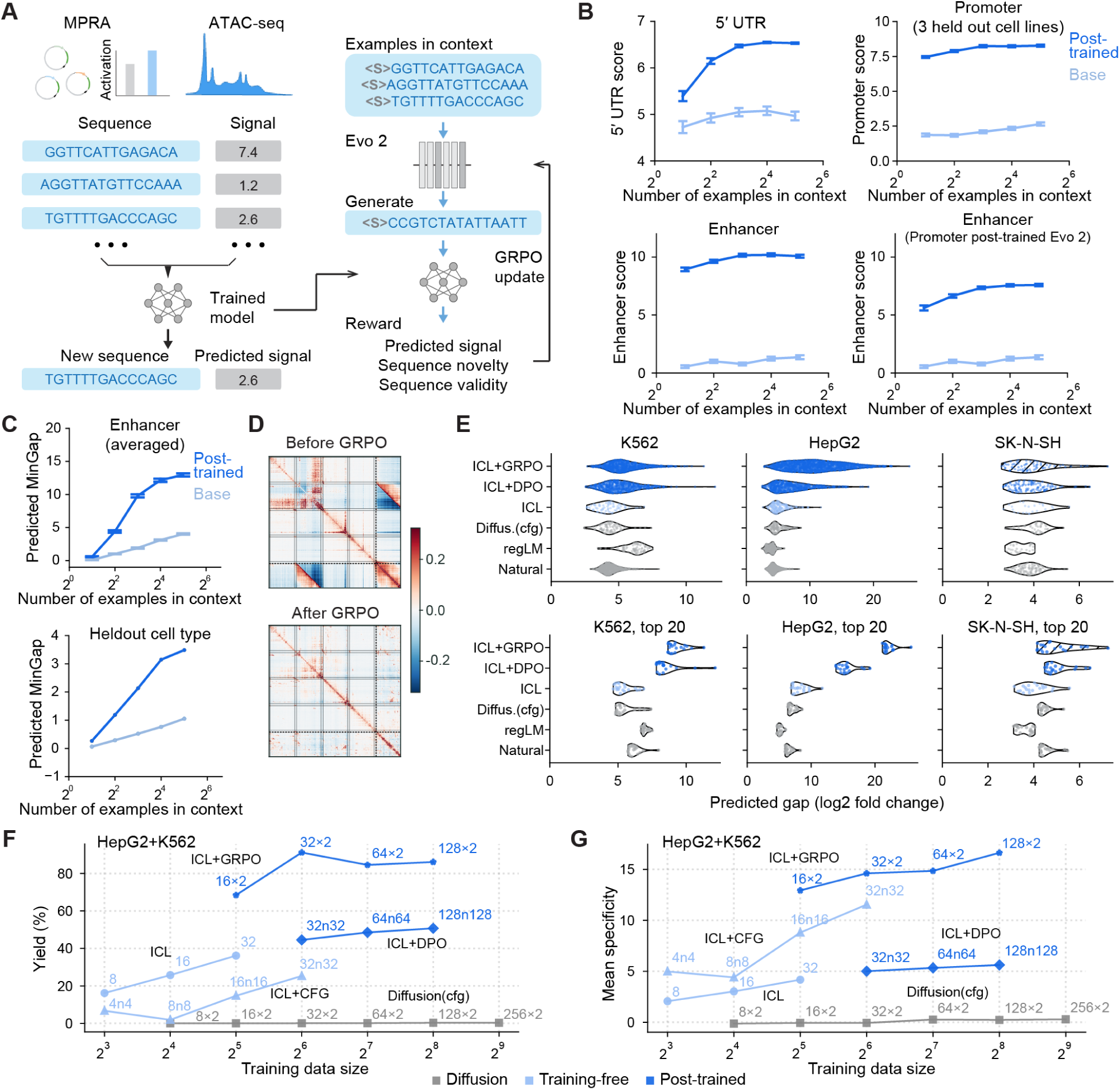
*|* **Incentivizing Evo 2’s in-context learning via preferences or virtual rewards. (A)** Predictive models trained on experimental measurements are used as virtual reward functions to reinforce few-shot sequence design with Evo 2. **(B)** Incentivizing Evo 2 1B with experiment-derived rewards increases the average predicted activity of generated sequences across multiple tasks: human 5*^′^* UTR design, human promoter design, and human enhancer design. For human promoter design, comparison was in three held-out cell lines. Post-training Evo 2 on the human promoter dataset also improves performance on human enhancer design across three non-overlapping cell lines. Error bars indicate SEM (n =32 prompts each condition). Light blue indicates base Evo 2, and blue indicates post-trained Evo 2. **(C)** Post-training Evo 2 7B with virtual rewards increases the average predicted specificity of generated enhancers. Evo 2 was rewarded for generating high-specificity sequences for HepG2 and K562 using 16 in-context examples, and was evaluated on test examples from HepG2 and K562, as well as the held-out cell line SK-N-SH. Top: comparison of average predicted specificity, measured by MinGap, across three cell types. Error bars indicate SEM (n =32 prompts each condition). Bottom: comparison of average predicted specificity in SK-N-SH. **(D)** Comparison of dependencies between generated enhancer sequences and in-context examples before and after reinforcement learning, measured using the categorical Jacobian. *(captioned continued on next page)* **(E)** Benchmarking Evo 2-based methods against established generative models for cell-type-specific enhancer design, with predicted specificity quantified in log2 fold-change. Evo 2-based methods include ICL with base Evo 2, ICL aligned by GRPO using experiment-derived rewards, and ICL aligned by DPO using experimentally measured sequences without an explicit reward function. All Evo 2-based methods use natural enhancers as in-context examples. Alternative approaches include DNA diffusion with CFG, trained on natural enhancer sequences; regLM, an autoregressive model fine-tuned on natural enhancers. As a reference, we also included natural enhancer sequences. Top panels: Predicted specificity of filtered sequences from a fixed number of generated candidates. Bottom panels: Predicted specificity of the top 20 sequences from each method. Colors denote method categories: post-trained Evo 2 ICL methods (blue), base Evo 2 ICL (light blue), and alternative methods or natural enhancers (gray). Hatching indicates that SK-N-SH was excluded from Evo 2 reinforcement post-training. **(F)** Data efficiency of different computational frameworks for cell-type-specific enhancer design, evaluated across different training dataset sizes using the synthetic enhancer dataset. Yield is defined as the proportion of generated sequences that are novel, valid, and specific to the target cell type. Training dataset size represents the total number of sequences used by each framework. The compared frameworks include DNA diffusion with classifier-free guidance (CFG; gray), training-free methods (light blue; ICL with base Evo 2 and Evo 2 ICL with CFG), and post-training methods (blue; Evo 2 ICL trained with DPO or GRPO). For ICL+GRPO, we leveraged the predictive model Malinois. For methods requiring both positive and negative examples (ICL+CFG and ICL+DPO), training dataset size includes both categories. For example, 32n32 denotes 32 positive and 32 negative examples (64 total). Similarly, for methods jointly trained on HepG2 and K562 enhancers (ICL+GRPO and DNA diffusion with CFG), dataset size represents the combined number of examples from both cell types. For example, 32*×*2 denotes 32 examples per cell type (64 total). For the training-free methods, i.e., ICL and ICL+CFG, the data size indicates the number of in-context examples. **(G)** Mean cell-type specificity of novel sequences generated by the computational frameworks evaluated in (**F**), across different training dataset sizes.

To test this, we first post-trained Evo 2 1B using group relative policy optimization (GRPO) and a predictor of 5*^′^* UTR activity (Shao et al., 2024; Zhang et al., 2025). The virtual reward combines normalized predicted activity with terms for DNA sequence validity, GC content, and sequence similarity (**Methods**). Using prompts sampled from a held-out test set, we found that reinforcement learning increased the predicted activity of generated 5*^′^* UTRs at every tested number of in-context examples (**Figure 2B**), indicating that predictive rewards can improve in-context example-conditioned generation.

We next applied this approach to human promoter and enhancer design. Using PARM, which predicts promoter activity across nine cell lines (Barbadilla-Martínez et al., 2026), we jointly post-trained Evo 2 on six cell lines and evaluated it on the remaining three cell types. After only tens of training steps, Evo 2-generated promoters showed significantly higher mean predicted activity in the held-out cell lines (**Figure 2B**). For enhancer design, we post-trained Evo 2 using Malinois, which predicts enhancer activity across K562, HepG2 and SK-N-SH human cell lines (Gosai et al., 2024). Using predicted activity in the target cell line as the functional reward increased mean predicted enhancer activity across all three lines (**Figure 2B**). These results suggest that post-training Evo 2 with experiment-derived virtual rewards is applicable across distinct regulatory design tasks. Interestingly, Evo 2 fine-tuned for human promoter activity also generated enhancers with improved predicted activity in HepG2, K562 and SK-N-SH, cell types which were not present during promoter reinforcement learning (**Figure 2B**).

Lastly, we post-trained Evo 2 7B using a reward that improves cell-type specificity in K562 and HepG2 measured by the gap between predicted activity in the target cell line and the highest predicted activity among off-target lines (MinGap as defined previously (Gosai et al., 2024)). We evaluated the resulting model in all three cell lines, including SK-N-SH, which was held out as a design target. Reinforcement learning increased predicted specificity in all three cell lines, including the held-out cell line SK-N-SH (**Figure 2C**). The improvement in SK-N-SH suggests that the model learned more general rules of cell-type information over trivial memorization.

To elucidate how reinforcement learning altered the influence of in-context examples, we compared categorical Jacobians before and after post-training. High-influence nucleotides were distributed more broadly across examples in the post-trained model than in base Evo 2 (**Figure 2D, S5A**). This pattern suggests that reinforcement learning promotes more effective integration of information across examples. We next asked how experiment-derived rewards reshape the biologically relevant statistics of generated sequences. Post-trained Evo 2 generated cell-type-specific enhancers with a higher density of transcription factor binding motifs enriched in the target cell type than base Evo 2 (**Figure S5B**). The motif density also scales with the number of prompted examples after post-training (**Figure S5B**). Moreover, replacing enhancer examples with random DNA sequences during post-training did not yield functional designs, indicating that post-training alone was insufficient without informative in-context examples (**Figure S5C**). These results demonstrate that aligning Evo 2 with experiment-derived rewards incentivizes it to extract and compose biologically relevant information more efficiently from in-context examples.

To overcome the limitation of using virtual reward models, we developed two approaches to enhance genomic ICL in Evo 2 without sequence-to-function models. First, we constructed two parallel prompts using celltype-specific and non-specific enhancer examples, and generated sequences based on the contrast signal from the two prompts, which we named classifier-free guidance (CFG). This training-free approach achieved better performance than genomic ICL with positive examples only (**Figure S6**). Second, we post-trained Evo 2 using direct preference optimization (DPO) to favor cell-type-specific over non-specific enhancers conditioned on shared ICL prompts (**Figure S7**) (Rafailov et al., 2023). Evo 2 post-trained via DPO also demonstrated higher performance on the enhancer design task than base Evo 2.

### Experimentally aligned Evo 2 designs highly specific enhancers for cell lines in silico

Cell-type specific enhancers have important applications in biological research and therapeutic engineering (Ben-Simon et al., 2025; Hunker et al., 2025). Designing such enhancers requires optimizing multiple objectives such as high activity in targeted cell types or states and low activity in off-target cells (de Almeida et al., 2024; Gosai et al., 2024; Sarkar et al., 2026; Taskiran et al., 2024). To achieve this design task, we applied genomic ICL with Evo 2 and further reinforcement learning, comparing against established design methods using either natural or synthetic enhancer datasets (Gosai et al., 2024).

Using the natural enhancer dataset, we compared MinGap specificity scores and sequence diversity of base and post-trained Evo 2, DNA-diffusion with classifier free guidance (CFG) (DaSilva et al., 2026) retrained on this dataset, and a specialized autoregressive DNA language model (regLM) (Lal et al., 2024). Base Evo 2 generated sequences with specificity broadly comparable to both baseline models without further parameter updates. Post-trained Evo 2 achieved higher specificity than both baseline models in two of three cell lines, while its top 20 designs achieved the highest specificity across all three cell lines (**Figure 2E**). Both base and post-trained Evo 2 maintained sequence diversity comparable to DNA diffusion and higher than regLM (**Figure S8B, C**). On the synthetic enhancer dataset, consisting of sequences designed with FastSeqProp (Gosai et al., 2024), post-trained Evo 2 achieved higher mean and top-20 predicted specificity than DNA diffusion with CFG retrained on this dataset, and improved over FastSeqProp for top-20 designs (**Figure S8A**). These results suggest that genomic ICL with base Evo 2 can be as effective as existing methods in designing cell-type specific enhancers, and ICL with post-trained Evo 2 obtained leading performance, overall designing sequences with higher quality and specificity.

To assess the data efficiency of ICL-based generation and post-training, we compared training-free Evo 2 ICL-based approaches and ICL post-trained using DPO or GRPO against DNA-diffusion with CFG across matched budgets of cell-type specific enhancers from the synthetic enhancer dataset (**Figure 2F, G, S9**). Without parameter updates, base Evo 2 ICL achieved yields of approximately 16–36% using only 8–32 in-context examples, whereas DNA diffusion produced negligible yields even with 512 training sequences (**Figure 2F**). ICL with CFG increased mean predicted specificity, although its yield remained below that of base ICL at matched sequence budgets (**Figure 2F, G**). Post-training of Evo 2 is also effective with limited labeled data. Using Malinois predictions as rewards, GRPO achieved yields of approximately 70% and 90% with total dataset budgets of 32 and 64 sequences, respectively, and obtained the highest mean predicted specificity among the evaluated methods, even on the held-out cell line (**Figure 2F, G, S9**). DPO achieved yields of approximately 45–50% with 64–256 positive and negative examples combined, although its mean predicted specificity was lower than that of GRPO. Evo 2 ICL therefore enables generation from small example sets at a level of data efficiency not matched by DNA diffusion under the tested conditions.

### Validating Evo 2-designed regulatory sequences in cultured cells

We then tested Evo 2-designed enhancers experimentally using a parallel reporter assay. For each cell line, we generated sequences with the base and post-trained Evo 2 models using natural or synthetic enhancer examples in context, and filtered them according to GC content and sequence novelty (**Figure 3A, 3B, S10A**). We included up to 40 designs per condition, and six natural and six synthetic enhancers with the highest specificity for each cell line as positive controls. We constructed a barcoded plasmid library in which each enhancer controlled the expression of an EGFP reporter through a minimal promoter, then sequenced the library to quantify enhancer–barcode abundance for each design (**Figure 3C**). We quantified the activation of each enhancer sequence by the log2 ratio of its corresponding normalized RNA to input plasmid DNA barcode abundance (log2 fold-change). Measurements from two independent transfections were highly correlated, with a mean Pearson correlation of 0.93 across the three cell lines, confirming the reproducibility of the assay (**Figure S10B**).

**Fig. 3.**
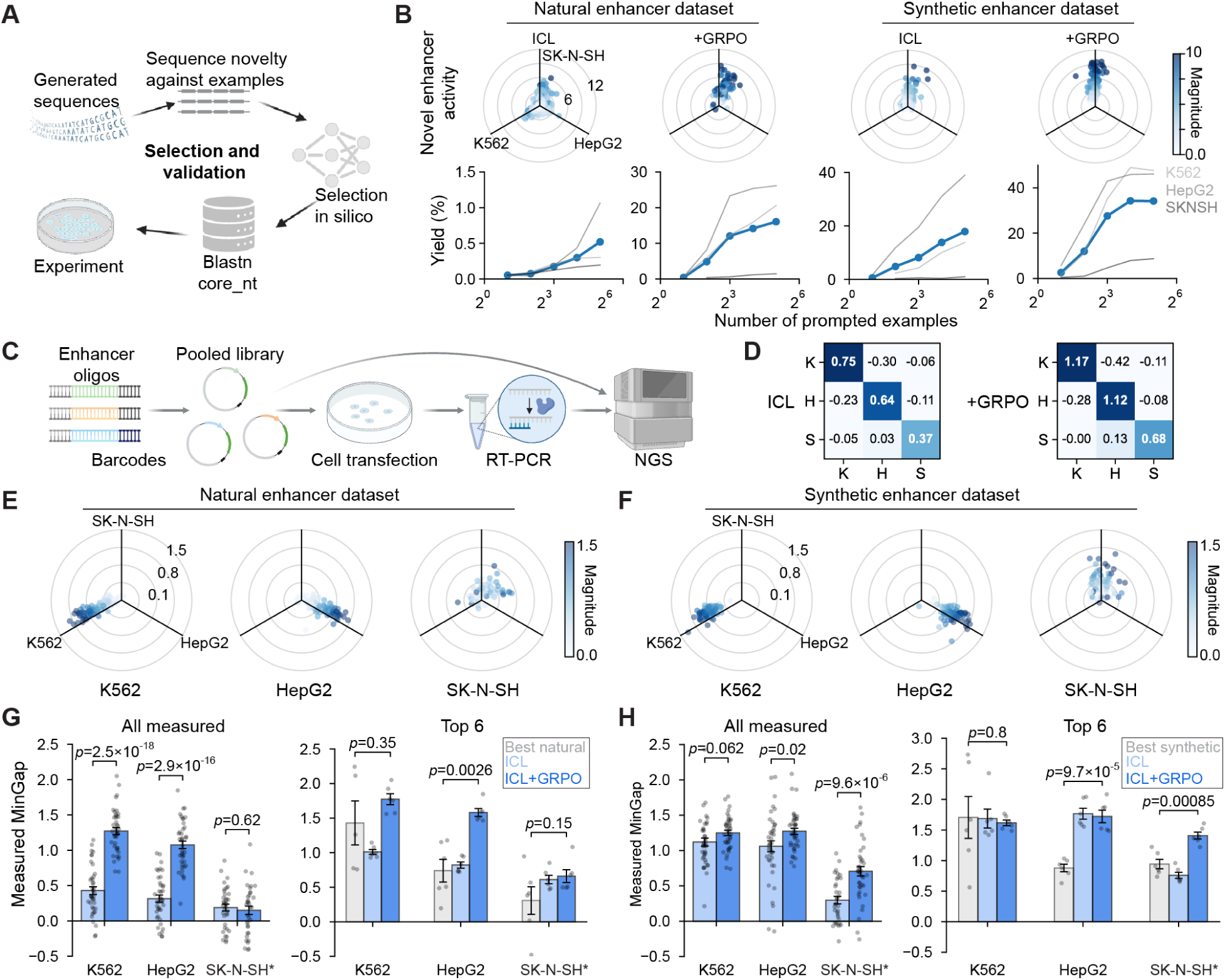
*|* **Experimental validation of cell-line-specific enhancers generated by Evo 2. (A)** Selection of enhancer sequences for *in vitro* validation. Generated sequences were filtered based on novelty relative to prompted examples, predicted specificity, and sequence novelty relative to the BLAST core_nt database. **(B)** Predicted activities and selection yields of Evo 2 generated enhancers in the held-out SK-N-SH cell line. Top panels: predicted activities of novel SK-N-SH enhancers in three cell lines before filtering for specificity, with each dot representing one sequence. Color indicates the total magnitude of predicted activity across three cell lines. From left to right: ICL with natural enhancer examples, ICL + GRPO with natural enhancers, ICL with synthetic enhancer examples, and ICL + GRPO with synthetic enhancers. Bottom panels: average yield of specificity-based selection across three cell lines. Gray lines represent yields for individual cell lines, ordered from light to dark: K562, HepG2, and SK-N-SH. **(C)** Experimental validation pipeline. Designed enhancers were synthesized with paired barcodes, cloned into plasmids containing EGFP, and delivered into cultured cells. After 48-hour expression, mRNAs were harvested, reverse-transcribed, amplified and sequenced. **(D)** Average measured activity in three cell lines for enhancers generated using base Evo 2 (left) or post-trained Evo 2 (right). The *x* -axis indicates measured activity in each cell line, and the *y* -axis indicates the target cell line. K, K562; H, HepG2; S, SK-N-SH. **(E)** Measured activity across three cell types for enhancers generated by ICL from natural enhancer examples. Each dot represents one generated sequence, and color indicates the magnitude of measured activity. From left to right: sequences designed for K562, HepG2, and the held-out SK-N-SH cell line. **(F)** Same as **(E)**, but for enhancers generated by ICL from synthetic enhancer examples. **(G)** Measured specificity of enhancers generated by ICL from natural enhancer examples across three cell types. Left: all measured enhancers targeting K562, HepG2, and the held-out SK-N-SH cell line. Right: top six enhancers for each target cell line. Gray, natural enhancers selected using a predictive model; light blue, ICL with base Evo 2; dark blue, ICL with post-trained Evo 2. *p*-values are from Student’s *t* -test. **(H)** Same as **(G)**, but for enhancers generated by ICL from synthetic enhancer examples.

Designed enhancers showed higher averaged activity in their target cell line than in off-target cell lines (**Figure 3D-F**). With natural enhancer prompts, post-trained Evo 2 generated sequences with higher measured specificity than the base model in K562 and HepG2 (**Figure 3G**), confirming that reinforcement learning directly improved Evo 2 for cell-type specificity. In the held-out cell type SK-N-SH, measured specificity of sequences generated by post-trained Evo 2 did not differ significantly from that of base Evo 2-generated sequences (**Figure 3G**). However, reinforcement learning increased the yield of sequences passing the predicted-specificity filter by 30-fold (**Figure 3B**). With synthetic enhancer examples as prompts, post-trained Evo 2 generated sequences with higher measured specificity than the base model in HepG2 and SK-N-SH, (**Figure 3H**), confirming post-trained Evo 2 could generalize when designing enhancers for a held-out cell-line target.

We also compared the top six designs of post-trained Evo 2 with the corresponding natural or synthetic positive controls and observed higher measured specificity in three of the six conditions, with no significant difference detected in the remaining three conditions (**Figure 3G,H**). These results show that Evo 2 designs can achieve specificity comparable to or higher than that of the top enhancers tested. Measured enhancer activities also correlated with Malinois predictions (**Figure S10C**, average Pearson correlation = 0.76), consistent with the published report (Gosai et al., 2024).

In summary, both base Evo 2 and post-trained Evo 2 generated enhancer sequences with desired cell-type specificity, and are comparable to or better than the best reported natural or synthetic enhancers. These experiments demonstrate that Evo 2 can generate functional, cell-type-specific enhancers via genomic ICL, and that reinforcement learning can increase the model’s in silico pass rates as well as improve measured specificity of ICL-generated enhancers.

### Fine-tuned AlphaGenome predicts chromatin accessibility across human brain cell types

We next sought to extend our design framework to designing enhancers targeting cell types *in vivo*. The brain’s remarkable cell-type diversity presents a challenging yet demanding setting for cell-type-specific enhancer design (Yao et al., 2023). Cell-type-resolved chromatin accessibility signals, such as single cell ATAC-seq, could provide a predictor for regulatory activities *in vivo* (de Almeida et al., 2024; Johansen et al., 2025; Kempynck et al., 2026; Taskiran et al., 2024). A cell-type specific enhancer can be characterized by a peak in ATAC-seq tracks, with high accessibility in the target cell type and low accessibility elsewhere. We therefore used a single-cell ATAC-seq dataset spanning 107 human brain cell types to develop predictive models for sequence evaluation (Li et al., 2023).

To establish a sequence-to-function map for cell-type specific enhancers, we fine-tuned AlphaGenome for two complementary tasks. First, we trained a classifier to distinguish candidate cis-regulatory elements (cCREs) from GC-matched genomic regions without cCRE annotations and random DNA sequences. The classifier (AG classifier) achieved an AUROC of 0.975 on a held-out test set (**Figure S11C**). Second, to determine the cell-type specificity of designed CREs, we fine-tuned AlphaGenome on approximately 500,000 cCREs to predict ATAC-seq signals across the 107 cell types. The resulting model (AG scorer) achieved a Pearson correlation of 0.73 between predicted and measured signals on a held-out test set (**Figure S11A,B**). To obtain orthogonal predictors aimed at detecting model-adversarial over-fitting in sequence design, we also fine-tuned Borzoi and Enformer (Avsec et al., 2021; Linder et al., 2025), which used different architectures and pretraining data, for classification and accessibility prediction. We combined four folds of Borzoi models with the Enformer model into an ensemble that exhibited similar performance as the AlphaGenome-based model (**Figure S11D-F**). This ensemble provided an independent mechanism for evaluating sequences generated using AlphaGenome-derived rewards.

### Post-trained Evo 2 designs highly specific enhancers for neuron types in vivo

We applied the reinforcement learning and ICL framework to design enhancers for neuron types *in vivo*. We selected four cell types as design targets *a priori* based on their morphological features and anatomical distributions. For each target, we identified 512-bp human genomic sequences corresponding to highly specific chromatin accessibility peaks in the targeted cell type and used them as in-context examples to prompt Evo 2. We split these sequences into training (60%), validation (20%) and test (20%) sets. Base Evo 2 generated novel sequences that were classified as cCREs with a yield of >25% (**Figure S12**). During reinforcement learning, we combined AlphaGenome-derived rewards, including a cCRE classification reward and a cell-type specificity reward, with sequence validity and novelty constraints, similar to the reward design for cultured cell lines (**Figure 4A**). After training, we sampled candidate enhancers by prompting Evo 2 with example sequences from the test set (**Figure 4A**).

**Fig. 4.**
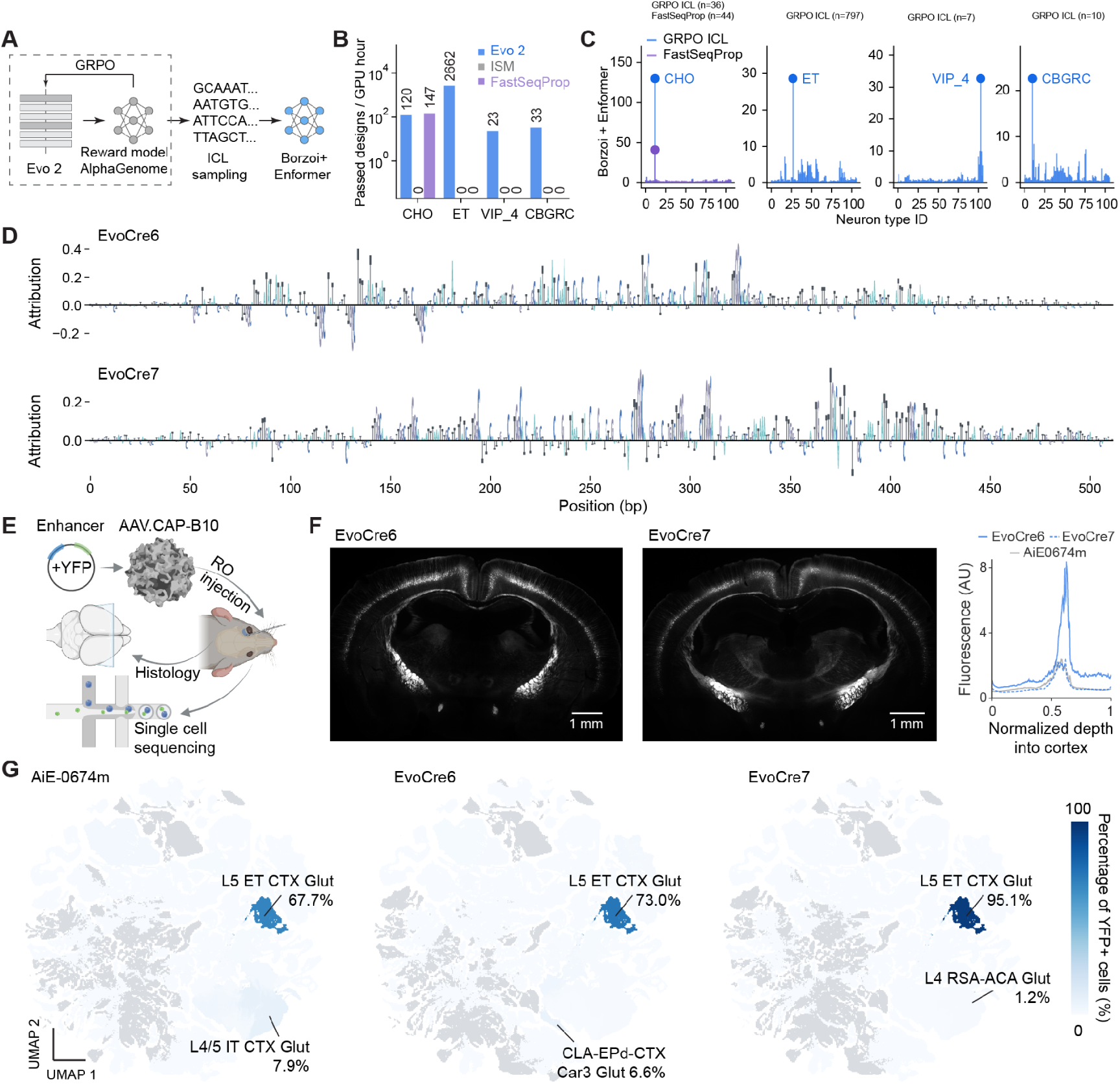
*|* **Designing and testing Evo 2-generated regulatory elements in the mouse brain *in vivo*. (A)** Design of enhancer sequences for *in vivo* validation. Evo 2 was post-trained using a reward model based on a finetuned version of AlphaGenome. Generated sequences were selected by a fine-tuned Borzoi+Enformer model ensemble. **(B)** Yield of filtering using the Borzoi+Enformer ensemble in four neuron types. ISM, in-silico mutagenesis. ISM and FastSeqProp uses the same reward function as Evo 2 post-training. **(C)** Average predicted activity for each of 107 neuron types in the human brain. **(D)** Attribution score at each position of Evo 2-generated enhancers. Two sequences are shown. **(E)** Workflow for testing Evo 2-generated enhancers in mice using enhancer AAVs. **(F)** Histology of two mice expressing designed enhancer AAVs for layer 5 extratelencephalic neurons, EvoCre6(left) and EvoCre7(middle). Right: fluorescence profile across cortex across three mice. Normalized depth indicates depth from the cortical surface normalized by the total depth of the cortex. Two mice were infected with designed enhancer AAVs. One was infected with AiE design as a positive control representing the current state-of-the-art enhancer. **(G)** Distribution of YFP+ cells across cell types for three tested enhancers targeting L5-ET. Cell types are mapped to the UMAP space of the Allen Mouse Brain Atlas. Top two cell types and their proportions are annotated. Gray indicates cell types below detection threshold in the data.

We filtered candidate sequences using the Borzoi–Enformer ensemble to identify and prevent reward hacking during reinforcement learning (**Figure 4A**). Post-trained Evo 2 generated designs that passed these filters for all four targets. Under the same AlphaGenome-based objective, *in-silico* mutagenesis (ISM) produced no designs that passed the Borzoi–Enformer filters, whereas a gradient based method, FastSeqProp (Linder and Seelig, 2021), produced passing designs for one target (**Figure 4B**). This suggests that ISM or Fast-SeqProp can overfit to the fine-tuned AlphaGenome models, whereas the additional constraints from Evo 2 ICL facilitate biological plausibility of the designed sequences that enables generalization to other scoring functions.

Selected Evo 2 designs showed high predicted chromatin accessibility in their intended target cell types and low background in other cell types (**Figure 4C**). We prioritized sequences with the highest predicted specificity and sequence novelty for experimental validations and refer to them as EvoCres. Attribution analysis of the Borzoi+Enformer ensemble’s specificity predictions highlighted central sequence regions containing clustered motifs (**Figure 4D**) (Sundararajan et al., 2017).

To test enhancer activity *in vivo*, we placed EvoCres or published enhancers that were mined from natural genomes upstream of a minimal promoter driving SYFP2, following the established enhancer-AAV design (Ben-Simon et al., 2025). We packaged the constructs into AAV.CAP-B10 vectors and delivered them to wild-type mice by retro-orbital injection (Goertsen et al., 2022). After three weeks of expression time, we collected and fixed brains for histology or single-cell sequencing (**Figure 4E**).

Seven of the nine EvoCres tested drove detectable expression in brain sections, and six produced spatially localized labeling, confirming that a substantial fraction of the selected designs were functional in the mouse brain. Specifically, EvoCre6 and EvoCre7 drove cortical labeling patterns consistent with the design target, layer 5 extratelencephalic (L5-ET) neurons. Their spatial expression patterns resembled those of a recently reported, natural L5-ET specific enhancer AiE0674m (**Figure 4F**) (Ben-Simon et al., 2025). Remarkably, at the same viral dose, EvoCre6 showed higher fluorescence than AiE0674m (**Figure 4F**). EvoCre9 and EvoCre11, which were designed to target Vip+ interneurons, drove expression in hippocampal neurons whose morphology is consistent with Vip+ interneurons (Turi et al., 2019), but also drove strong expression in hippocampal CA3 (**Figure S13**).

To quantify cell-type specificity, we next characterized the molecular identity of the YFP+ cells by wholebrain single-cell RNA sequencing and cell-type classification using MapMyCells (Daniel et al., 2026). Strikingly, 73% and 95% of YFP+ cells were assigned to the L5-ET neuron type for EvoCre6 and EvoCre7, respectively, compared with 68% for AiE0674m (**Figure 4G**). Across different thresholds for defining YFP+ cells, EvoCre7 showed consistently better specificity while EvoCre6 showed similar specificity to AiE0674m (**Figure S14**). For EvoCre9 and EvoCre11, 58% and 74% of YFP+ cells are classified as CA3 glutamatergic neurons, respectively. These designs therefore primarily labeled a population different from the design target. Nevertheless, their targeting profile suggests potential utility for labeling CA3 excitatory neurons. We hypothesize that these misspecified or failed designs could be due to differences in the regulatory landscape between the human and the mouse brain, an issue that could be addressed by jointly filtering using predictive models trained on mouse and human ATAC-seq datasets. Taken together, these results establish the feasibility of using enhanced in-context learning that leverages human chromatin accessibility data to generate functional enhancers for targeting neuron types *in vivo*.

## Discussion

In this study, we demonstrate that generative in-context learning (ICL) emerges in Evo 2, enabling the design of novel functional non-coding sequences without further training or steering. Evaluated by independently trained predictive models, Evo 2 generated functional sequences spanning five regulatory modalities with 30-50% yield and <60% sequence similarity from only 4-64 in-context examples. We also observed in-context scaling in which more example sequences in the prompt led to better generated sequences. Mechanistically, we observed a few attention modules that are causally linked to the inductive behavior and ICL of Evo 2. Our findings establish genomic ICL as a novel framework for sequence design using a general-purpose generative genome model that can work in a low-data regime.

The ability to generate novel sequences based on in-context examples can be further strengthened by virtual rewards, including large sequence-to-function models or simple statistical properties such as GC content. This is analogous to aligning generative language models by reinforcement learning from human feedback or with virtual reward models (Ouyang et al., 2022), which has led to strong improvements in other domains. One concern of this framework is reward hacking, in which the model generates sequences with strong *in-silico* scores but weak biological plausibility. Our results suggest that incentivizing Evo 2 using a virtual reward can be effective, although the correlation between measured and predicted activity can still be improved.

While generative ICL does not strictly require a sequence-to-function model, better predictive models (Avsec et al., 2021, 2026; Chen et al., 2025b; Kempynck et al., 2026; Lal et al., 2025, 2026; Linder et al., 2025; Pampari et al., 2025; Zhou, 2022) are highly complementary and will directly benefit both reinforcement-learningbased post-training of Evo 2 ICL and downstream filtering. Importantly, the information learned by Evo 2 provides additional regularization during sequence optimization; indeed, we observed that black-box in-silico mutagenesis or gradient-based optimization of sequence-to-function models struggles to produce designs that pass orthogonal filters compared to reinforced Evo 2 ICL (**Figure 4B**). These results mirror findings from protein design in which structure- or function-conditioned sequence design models help prevent adversarial generation (Brookes et al., 2019; Dauparas et al., 2022).

The effectiveness of ICL depends on the information available in its examples. When examples are extremely scarce, ICL may favor directly copying the prompted examples instead of generating novel sequences. Optimizing prompt composition or incorporating functionally relevant homologous sequences could enrich the context and improve generation. Although ICL reduces the need for task-specific generators, our reinforcement learning approach, GRPO, relies on a predictive function to align Evo 2 to task-specific knowledge. When such models are unavailable, classifier-free guidance or preference-based post-training methods such as DPO can still improve ICL performance using contrasting examples or experimentally measured preferences, albeit with a potential decrease in design success rates (**Figure 2F,G**). Finally, our causal perturbations identify components that contribute to ICL but do not fully resolve its underlying mechanism. Further work is needed to determine how these components interact and whether ICL in Evo 2 depends on distributed and redundant circuits.

The methodology is readily generalizable to designing enhancer sequences that control gene expression in diverse cell types in mammalian systems *in vivo*. In our study, we demonstrated this by designing and testing Evo 2-generated regulatory sequences in the mouse brain and achieved 2 out of 2 successes in one neuronal subtype. Recent publications have demonstrated synthetic enhancer design in insects (de Almeida et al., 2024; Taskiran et al., 2024) and vertebrates (Kempynck et al., 2026), as well as tissue-specific enhancer design in mouse embryos (Chen et al., 2025a). Our work extends these advancements to finer resolution in the mammalian brain by optimizing predicted enhancer specificity for single neuronal types against 106 other neuron types. Notably, the sequences for fine-tuning sequence-to-function models and prompt Evo 2 are all from the human genome, highlighting shared regulatory codes across species captured by Evo 2 as a sequence generator.

Evo 2 was trained on a large corpus of natural genomic sequences and ICL emerges potentially because of data and model scaling. ICL could transform pre-trained genomic language models into universal generators of biological sequences, including DNA, RNA, proteins and even whole genomes (Brixi et al., 2026a). Looking forward, future work could enhance this meta-learning behavior during model training and evaluate generalization to broader biological tasks with scale. While we have only shown results for sequence generation, we also aim to explore the possibility of leveraging ICL in Evo 2 in predictive tasks in data-scarce regimes. Leveraging and enhancing ICL in generative genomic models could accelerate the design and discovery of complex biological systems.

## Methods

### 5′ UŁR generation and in-silico evaluation

Functional 5*^′^* UTR sequences were taken from a published MPRA dataset where 280,000 oligos were measured for mean ribosome load (MRL) (Sample et al., 2019). We filtered the measured 5*^′^* UTRs according to MRL with a threshold of 8.0 and selected active sequences for prompt construction. Prompts with 4, 8, 16, 32, 64, 128 or 256 examples (50 bp each) were used to condition Evo 2 7B to generate novel 5*^′^* UTRs with nucleus sampling (temperature 1.0, top-k 4, top-p 0.9). Generated sequences were first filtered using basic quality filters (GC content below 65%) and novelty filters (pairwise sequence identity below 0.6 and local blastn alignment score below 30 against every prompt sequence), and evaluated using a predictive PRED-5UTR model (Zhang et al., 2025).

### IRES generation and in-silico evaluation

Functional IRES sequences were drawn from a published high-throughput screen in which a *∼* 55,000-oligo library was assayed for cap-independent translation activity using a bicistronic luciferase reporter (Weingarten-Gabbay et al., 2016). We restricted the library to the 8,940 human-genome-derived sequences and ranked them by eGFP measurement, selecting the top-scoring sequences for prompt construction. Prompts comprising 2, 4, 8, 16, or 32 examples, each 174 bp in length, were used to condition Evo 2 7B to generate novel human IRES elements with nucleus sampling (temperature 1.0, top-k 4, top-p 0.9). Generated sequences were filtered with basic quality filters (GC content below 70%) and a novelty filter (pairwise sequence identity below 0.6 and local blastn alignment score below 60 against every prompt sequence), and scored in silico with DeepIRES (Zhao et al., 2024), a sequence-to-function model employing a convolution and attention-based architecture.

### Human promoter generation and in-silico evaluation

Human promoter sequences were taken from a published MPRA study where about 4 million genomic sequences were measured across 9 cultured cell lines, namely K562, HepG2, HEK293, HAP1, AGS, HCT116, U2OS, MCF7, and LNCaP (Barbadilla-Martínez et al., 2026). We took the sequences that are 200-230 bp taken from the positive strand. We selected sequences with top measured and predicted activities. For each transcription starting site (TSS), we selected one promoter to prevent overlapped sequences. For each of the 9 cell types, we prompted Evo 2 7B with 2, 4, 8, 16, 64 or 128 sequences flanked by a random separator sequence (‘ATGCATGA’, the default separator) to generate novel human promoters with nucleus sampling (temperature 1.0, top-k 4, top-p 0.9). Generated sequences (230 bp each) were filtered using basic quality filters (GC content between 25% and 75%, no separator sequence in the middle of the promoter, and longest homopolymers is shorter than 8-mer) and novelty filters (pairwise sequence identity below 0.6 and local blastn alignment score below 60 against every prompt sequence), and evaluated using the corresponding PARM model for the cell type (Barbadilla-Martínez et al., 2026). We have empirically found that using a single ‘N’ as the separator yielded sequences with similar predicted activities, and caused higher chance of generating intact sequences without emitting premature separator in the middle of generations.

### Human enhancer generation and in-silico evaluation

Human enhancer sequences for three cell types (HepG2, K562 and SK-N-SH) were taken from a published MPRA study (Gosai et al., 2024). We took both 200 bp natural genomic sequences (referred to as natural enhancer dataset) or synthetic enhancers designed using FastSeqProp (referred to as synthetic enhancer dataset) that are measured to be highly active. For each of the three cell types, we prompted Evo 2 7B with 2, 4, 8, 16, 32 or 64 sequences to generate novel human enhancers with nucleus sampling (temperature 1.0, top-k 4, top-p 0.9). Generated 200 bp sequences were filtered using basic quality filters (GC content between 25% and 65%, no separator sequence in the middle of the enhancer, and longest homopolymers is shorter than 8-mer) and novelty filters (pairwise sequence identity below 0.6 and local blastn alignment score below 60 against every prompt sequence), and evaluated using the Malinois model for the cell type (Gosai et al., 2024).

### Yeast promoter generation and in-silico evaluation

Yeast promoter sequences (80 bp each) were taken from a published dataset (Vaishnav et al., 2022). We first deduplicated sequences based on first and last 6 nucleotides, and selected the sequences that are measured to be highly active. We then prompted Evo 2 7B with 1, 2, 4, 8, 16, 32 or 64 sequences to generate novel yeast promoters with nucleus sampling (temperature 1.0, top-k 4, top-p 0.9). Generated 80 bp sequences were filtered using basic quality filters (GC content below 65%) and novelty filters (pairwise sequence identity below 0.6 and local blastn alignment score below 50 against every prompt sequence), and evaluated using the regLM model (Lal et al., 2024).

### Identification and ablation of inductive heads in Evo 2

We localized attention heads underlying induction in Evo 2 models using a two-step screen applied to every attention head (60 heads in Evo 2 1B, 160 heads in Evo 2 7B). First, we identify a list of attention heads that use repeated patterns in the prompt to predict next tokens. To do this, we constructed 100 self-concatenated random sequences *S* +*S*, where *S* was 64-bp random DNA sequences. We computed the mean attention scores between corresponding positions in the tandem sequences. Second, we computed an ‘induction head score’ to identify attention heads that recognize the ICL task structure. We constructed ICL prompt by using 8-shot HepG2 enhancer sequences flanked by separating sequences. The attention head score is average attention from the final position of each separator to the token immediately following every preceding separator. We took the attention heads with high induction head scores from each layer for ablation studies.

To evaluate the effects of ablating an induction head, we prompted the model with n-shot HepG2 enhancers (n=2, 4, 8, 16, 32) sampled from the synthetic enhancer dataset. For each n, we constructed 32 prompts and sampled 8 sequences each prompt. In the Evo 2 1B model, we ablated the induction head L3H6 (layer 3 head 6), or a random head L3H7 of the same layer as a control. In the Evo2 7B model, we ablated two induction heads L3H21 and L10H17. As a control, we ablated L3H22, and L10H20. To ablate an attention head, we set the output matrix corresponding to the attention head to be 0, instead of zeroing the attention scores directly. Generated sequences were evaluated using Malinois (Gosai et al., 2024).

### Sparse autoencoder feature analysis

For each of three cell lines (HepG2, K562, and SK-N-SH), a 32-shot enhancer ICL prompt was concatenated with a corresponding generated sequence, and layer-26 activations from *evo*2_7*b*_262*k* were encoded using the mixed eukaryotic-prokaryotic SAE from Brixi et al (Brixi et al., 2026b). For each SAE feature, mean activation trajectories were calculated across either the 32 example bodies or the 33 contiguous separator regions, averaged across cell lines, and correlated with block index. Only features active in approximately half or more of the corresponding regions were considered. The four most strongly correlated features under each pooling scheme were retained and visualized.

### Categorical Jacobian

We analyzed model generations from Evo 2 7B for HepG2, K562, and SK-N-SH before and after GRPO using the Categorical Jacobian calculation for prompts with 4, 8, and 16 examples (Zhang et al., 2024), selecting one generation from each of 16 prompts, for a total of 288 maps. For a sequence of length *L*, each position *i* underwent complete mutagenesis with substitutions A/C/G/T. Here, *L* covers the length of the concatenated prompt, separator sequences, and generation. For each substitution at each site, we then computed the change in the model’s output logits for every subsequent position j. This led to a 4D tensor *J_i,a,j,b_* where *i*, *j* are sequence indices and *a, b* are the substituted and output-logit nucleotides respectively. We mean-centered and symmetrized this tensor by averaging entries with their input-output transposes, then reduced this to a raw *L × L* contact map by computing the L_2_ norm over nucleotide pairs:

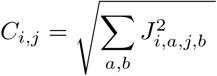

and set the diagonal to zero. We also computed an average-product-corrected map using the diagonal-zero map *C* for visualization, again setting the diagonal to zero:

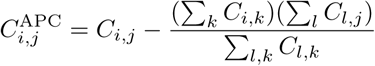

For visualization, we cap the color scale at the 99^th^ percentile of absolute values pooled from both APC maps.

### Example influence analysis

Using the Categorical Jacobians from above, we then analyzed how strongly the model’s output logits are sensitive to individual examples in the prompt, and how concentrated this influence is.

First, we ask whether one examples dominates the model’s contacts for the generated sequence. We add all example-to-generation contacts, find the support with the largest share, and normalize for the number of sequences in context. Specifically, for each prompted base, we summed its raw contacts with all 200 generated tokens and let *M* be the sum of these contact values for examples *i* and there are *k* examples in-context, and let *p_i_* = *M_i_/* ∑*_k_ M_k_*.We report the metric

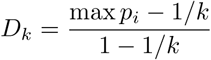

which is zero when all examples contribute equally to the generation and one when one example sequence accounts for all example-generation contacts.

Second, we ask how many “effective examples” there are by checking how many examples contribute to the 40 bases with the largest summed contacts to the generated sequence. Among those 40 selected bases in the prompt, we let *n*_i_ be the number of those bases belonging to example *i*, and compute

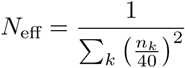

which is one when all 40 selected bases come from one example, and is maximized when the 40 selected bases are distributed as evenly as possible across examples. For each (cell type, pre/post-GRPO, number of examples in context), we calculated the metric of interest once for each of 16 prompt-generation pairs. Error bars show 95% bootstrap intervals for the mean of that metric, obtained by resampling those 16 pairs with replacement 4,000 times.

### Motif Enrichment Analysis

We used the 1,019 position frequency matrices (PFMs) in the JASPAR 2026 CORE vertebrate non-redundant PFM collection (Ovek Baydar et al., 2026). For each PFM position, we added a 0.5 pseudocount to each A/C/G/T count, divided by the column total plus 2, and converted the resulting probabilities to log2 odds against a uniform background to obtain position weight matrices (PWMs). We then scanned each sequence and its reverse complement by summing these weights over every possible window, counting a window as a hit if its score was at least *S* _min_+0.9(*S* _max_ – *S* _min_), where *S* _min_, *S* _max_ are the sums of the position-wise minimum and maximum weights based on the PWM. A forward and reverse-strand hit for the same PWM at the same position counted as one hit, but overlapping starts or matches to different PWMs counted separately.

We calibrated our set of cell-enriched matrices on natural data. We applied the same scanning procedure to 197 HepG2, 1,266 K562, and 85 SK-N-SH sequences. Each sequence was called a “carrier” for a PWM if it contained at least one hit, and a PWM was assigned to a cell type if a cell type had at least 10 carriers, at least 10% of sequences in that cell type were carriers, and that cell type’s carrier prevalence was at least 5 percentage points higher than in the other cell types. We address GC-content confounding by running a one-sided Cochran-Mantel-Haenszel test to check for enrichment against the other two cell types within 5%-wide GC-content bins, requiring that at least 80% of the sequences in each cell type fell into overlapping GC-content bins for comparison. Afterwards, we adjusted the resulting p-values afterwards using the Benjamini-Hochberg procedure, considering BH-adjusted q*≤*0.05 as significant. This led to 75 HepG2, 25 K562, and 50 SK-N-SH enriched matrices.

For every Evo 2-generated sequence, we counted hits to matrices enriched in its target cell type and hits to matrices enriched in off-target cell types. We analyzed eight generations from each of 16 matched prompts per condition for a total of 2,304 sequences. We compute the same counts for the natural-reference sequences used to define the cell-enriched PWMs, and separate natural sequences predicted to be inactive in each target cell type by Malinois for comparison.

### Incentivizing Evo 2 with experiment-derived virtual rewards

We fine-tuned Evo 2 with group relative policy optimization (GRPO) (Guo et al., 2025; Shao et al., 2024), a critic-free policy-gradient method, based on Evo 2 ICL inference to improve the quality of generated sequences. In each training step, we sampled n-shot examples (n=16 by default) from the training dataset and concatenated them to prompt Evo 2 to generate *G* designs. For each design *o*_i_, we assign it a reward score *R*(*o*_i_). The rewards can include rule-based sequence-validity and formatting terms, including using A, T, C, G for DNA sequences only (weight=0.2), no separator emitted in the middle of generated sequences (weight=0.3), proper GC content (weight=0.5), and sequence novelty (low sequence identity to in-context examples, weight=1.0). Rewards also include task-specific terms such as predicted activity or specificity (weight=1.0). We then updated the model by gradients computed using the following GRPO objective

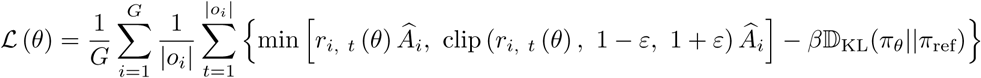

where the group-relative advantage is

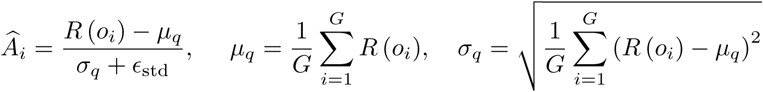

and the per-token importance ratio is

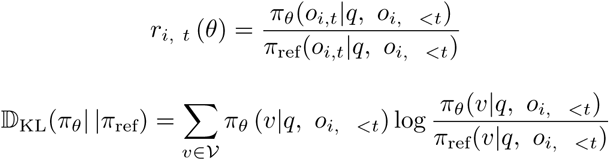

Here the KL distance to a frozen reference model regularizes drift during training. We optimize the model with AdamW at a learning rate of 10*^−^*^6^. We evaluated the model periodically by prompting the model with examples from the validation dataset and compute the rewards on generated sequences, and selected best checkpoints based on the mean validation reward. For enhancer or promoter generation for cultured cell lines, we cycled through cell lines during training, with each cell type carrying its own task-specific rewards. For Figure 2B, we post-trained the Evo 2 1B model. We use PRED-5UTR model (Zhang et al., 2025) (clipped to [2.0, 8.0] and normalized to [0,1]) to provide task-specific reward for 5*^′^* UTR, PARM model (Barbadilla-Martínez et al., 2026) (clipped to [0.0, 5.0] and normalized to [0,1]) to provide task-specific reward for human promoter, and Malinois model (Gosai et al., 2024) (clipped to [0.0, 6.0] and normalized to [0,1]) to provide task-specific reward for human enhancers. For improving enhancer specificity in Figure 2C, we post-trained the Evo 2 7B model. We used predicted MinGap by Malinois model (Gosai et al., 2024) (clipped to [0.0, 3.0] and normalized to [0,1]) as the task-specific reward, and held out SK-N-SH as design target.

### Classifier free guided generation using Evo 2

We implemented classifier free guidance (CFG) as contrastive decoding over two sets of in-context examples. At each decoding step, two next-token logit vectors were computed in parallel using the two prompts, and then were combined as *logits*_guided_ = *logits*_cond_ + *w×*( *logits*_cond_ *− logits*_base_). Here *logits*_cond_ were computed from the prompt with positive examples that one wants to steer towards, and *logits*_base_ were computed with baseline or negative examples. The two tracks keep separate caches but are advanced by the same sampled token, so the contrast stays aligned across the full generation. Baseline or negative examples were a shared pool of non-specific enhancers. We swept the steering strength *w* over [0.5, 1, 2]. Decoding temperature T was 1.0 by default. We also swept T over [0.75, 1.0, 1.25] to understand how T and *w* jointly influence the quality of generated sequences. Sampled sequences were evaluated with: (1) sequence novelty, as measured by sequence similarity with prompted examples (both blastn local alignment scores and global sequence identity), and (2) enhancer specificity, as predicted by the Malinois model (Gosai et al., 2024).

### Direct preference optimization

To demonstrate that we can also improve Evo 2 generation without external reward function, we aligned Evo 2 for ICL-steered enhancer design using direct preference optimization (DPO) (Rafailov et al., 2023). For each DPO step, we constructed a triplet (*x*, *y* _w_, *y* _l_) where *x* is the shared prompt containing 16 enhancer examples, and *y* _w_ is a cell-type specific enhancer (a preferred response), and *y* _l_ is an enhancer without desired cell-type specificity (a negative response). The log-likelihood of generated responses was computed as log *π*(*y* |*x* ) = Σ_t_ log *π*(*y* _t_ | *x*, *y* <*t* ). The policy *π_θ_* (i.e. improved Evo 2) was optimized against the base model *π*_ref_ (i.e. Evo 2) under the standard DPO objective with *β* = 0.1.

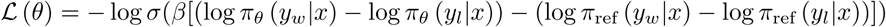

We optimized the objective used AdamW at a constant learning rate of 5 *×* 10*^−^*^7^. HepG2 and K562 were interleaved during training so both cell types were co-trained within a run.

### DNA-diffusion model with classifier free guidance

We trained DNA-Diffusion models (with classifier-free guidance, CFG) using the training recipe in (https://github.com/pinellolab/DNA-Diffusion) (DaSilva et al., 2026). To achieve CFG, cell-type labels were replaced by a reserved unconditional token with probability p_uncond_ = 0.1 during training. Both models were trained independently on a single NVIDIA H100 GPU using fused Adam (learning rate 4 *×* 10*^−^*^4^). We trained models with early stopping if the validation loss saturated. For the natural enhancer dataset (10941 total), we trained the model on 9525 enhancers and validated using 734 enhancers. We also trained the model on highly specific natural enhancers only (1592 total), but the sampled sequences were highly degenerated (Figure S8B). For the synthetic enhancer dataset (3304 enhancers total), we trained the model on 1981 highly specific enhancers and validated using 659 enhancers. The best checkpoint by validation loss was selected for each model. After training, we sampled 1000 sequences per cell type from each trained model by running the reverse diffusion process conditioned on each cell-type label using CFG with a guidance strength of 1.0 and a sampling batch size of 10. Sampled sequences were filtered by Malinois prediction (on-target activity>4.0 and off-target activity<1.6) and evaluated for predicted MinGap in Figure 2E and S8A. The same activity filter was applied to sequences from other methods too.

### Autoregressive enhancer model

We directly took sampled enhancer sequences (Table S6 from Lal et al (Lal et al., 2024)) from regLM and filtered them by Malinois prediction (on-target activity>4.0 and off-target activity<1.6) and evaluated them for predicted MinGap in Figure 2E and S8A.

### Benchmarking data efficiency in generative enhancer design methods

We compared the performance and data efficiency of different generative frameworks for cell-type-specific enhancer design across varying training dataset sizes. For methods requiring training or fine-tuning, we restricted training to HepG2 and K562 enhancers, holding out SK-N-SH as an unseen design target. We constructed nested training subsets of progressively doubling sizes from a single random shuffle of positive examples, such that each subset of size *N* was contained within the subset of size 2*N*. For methods requiring negative examples, we independently sampled *N* negative examples to supplement *N* positive examples. Dataset size refers to the total number of sequences used by each method, including positive and negative examples, or examples from two cell types. For example, a dataset with 32 positive and 32 negative examples, or 32 examples each from HepG2 and K562 was labeled with a size of 64. For methods requiring training, datasets were further split into 80% training and 20% validation. Training-free Evo 2 ICL methods were limited to 32 in-context examples due to the 8,192-token context window, while Evo 2 post-training used 8 or 16 in-context examples per prompt. Model training and sequence generation followed the procedures described in the corresponding Methods sections.

For each (method, dataset size, target cell type) condition, we generated 1,024 sequences and evaluated them using a shared pipeline. Three rule-based filters were applied: (1) DNA validity, i.e. exactly 200 bp containing only A/C/G/T; (2) GC content, requiring a range between 0.30 and 0.70; and (3) design novelty, requiring a maximum blastn local-alignment bit score below 60 against the corresponding training pool, considering both forward and reverse-complement orientations. Sequences passing these filters were scored using Malinois to predict cell-type-specific enhancer activity. A sequence was classified as specific if its predicted on-target score exceeded 4.0 and its maximum off-target score was below 1.4. Design yield was defined as the fraction of all generated sequences satisfying all four criteria. Mean specificity was quantified by MinGap, defined as the difference between predicted on-target activity and maximum off-target activity, averaged across all sequences passing the three rule-based filters. Results for HepG2 and K562 are shown in Figure 2, and those for the held-out SK-N-SH cell type in Figure S9.

### Łraining and evaluating enhancer classifiers

We added a logistic regression head (3073 parameters) to the AlphaGenome trunk with 128-bp resolution and no paired embeddings to construct the classifier model. In the enhancer classification dataset, positive class includes 525,473 unique cCREs taken from the CATlas human brain atlas (restricted to autosomes and filtered against the ENCODE hg38 v2 blacklist with a 50 bp window (Amemiya et al., 2019)), while the negative class includes 80% of GC-matched genomic background and 20% of uniform random sequences. Train/validation/test splits were by chromosome: chr10 for validation, chr11 for test, and the remaining for training. Training sequences were augmented with random shifts of up to *±*32 bp within the 576 bp window. Binary cross-entropy was used to train the model. We used Adam with learning rate 10*^−^*^4^ and trained it on one NVIDIA H100 for one epoch. The model was then evaluated on the held-out chr11 test set (62,965 sequences). The training infrastructure was modified based on Lal et al (Lal et al., 2025) and Kempynck et al (Kempynck et al., 2026). We also trained classifiers based on the Borzoi trunk (4 folds) and the Enformer trunk with the same setting, to be independent models compared to the AlphaGenome-based model.

### Łraining and evaluating enhancer scorers

The scorer predicts cell-type-resolved chromatin accessibility for 512-bp cCREs. We added a convolutional head (328,811 parameters) to the AlphaGenome trunk with 128-bp resolution and no paired embeddings to construct the classifier model. Because 512 bp input is short relative to the trunk’s receptive field, we tiled input sequences twice along the length axis. We trained the model on the human CATlas brain cCRE set (525,293 regions) with a chromosome-level data split (chr10 for validation, chr11 for test, and the remaining for training). Training labels were average ATAC-seq signals across 107 cell types scaled by a factor of 0.1. High-variance cCREs (62,126 regions selected for high variance across cell types) were upweighted by 12-folds during training and we randomly shuffled the training data. We optimized the model with a Poisson negative log-likelihood loss using Adam at a learning rate of 10*^−^*^4^ on a single NVIDIA H100. The best checkpoint was evaluated on all 27,264 held-out chr11 cCREs in the label_scale = 10 target space. Averaged per-cell-type Pearson r was 0.7264 (median 0.7291, range 0.549–0.817) and the pooled correlation over all predictions on the test set was 0.7332. We also trained scorers based on the Borzoi trunk (4 folds) and the Enformer trunk with the same setting.

### Enhancer generation and selection for in vivo validation

For generating enhancers for neuron types *in vivo*, we post-trained Evo 2 7B with GRPO, as we previously described. We trained an independent policy network for each of four CATlas brain cell types (CBGRC, CHO, ET-L5, and VIP_4). For each cell type, curated 512 bp cCREs with desired cell-type specific ATAC peaks were partitioned 60%/20%/20% into training, validation and test sets (153/51/52 sequences). We used 16-shot example prompting to generate rollouts by nucleus sampling (temperature 1.0, top-*k* 4, top-*p* 0.9). We assign each rollout sequence with a scalar reward with a maximum of 4.0, formed as a weighted sum of six terms: (i) sequence validity (weight 0.2, reward=1 if the rollout contained only uppercase A/C/G/T); (ii) no separator in the middle (weight 0.3, reward=1 if the separator did not occur in the middle of the rollouts, i.e. no premature termination of sequence generation); (iii) GC content (weight 0.5, reward=1 if GC content is in the range of [0.25, 0.65]); (iv) sequence novelty (weight 1.0, reward=1 if the maximum BLASTN-scored pairwise sequence identity to the in-context examples was *≤* 60%, and 1*−*identity otherwise); (v) a CRE confidence term (weight 1.0, reward=min(*p*/0.8, 1) for predicted cCRE probability *p*); and (vi) a cell-type specificity term (weight 1.0) computed from a 107-cell-type chromatin-accessibility model as clip(*g* /3.6, 0, 1). Here *g* is the stronger MinGap computed as *g* =*A*_on_*− k·*max(0, max(*A*_off_) *− τ* ) to constrain off-target activity *A*_off_ to *τ* , while prioritizing the on-target activity *A*_on_. We optimized the model using the GRPO loss described before, where we set the importance ratio clip *ε*=0.2 and the KL coefficient *β*=0.005. Training proceeded in two stages of 600 steps: stage 1 initialized from the base model with a specificity shape (*τ* =1.2, *k* =2.0) at learning rate 1 *×* 10*^−^*^5^, and stage 2 resumed from the best stage-1 checkpoint with a specificity shape (*τ* =0.6, *k* =4.0) at learning rate 5 *×* 10*^−^*^6^. Final sequence sets were generated from these checkpoints using prompts built from the held-out test split at 8, 16 and 32 in-context examples, with 128 prompts *×* 8 completions = 1,024 sequences per cell type and shot count.

### Molecular cloning

To construct the MPRA library, we ordered 276 bp oligos from Twist Bioscience that contain 200 bp enhancers, 28 bp connecting sequences, 15 bp barcodes, and adapter sequences on either end. We amplified the oligos for 12 cycles by PCR and inserted the amplicon into a pcDNA3.1 vector, then transformed them into Escherichia coli (XL10-Gold, Agilent Technologies). We estimated 200-300 colony-forming units (CFUs) per oligonucleotide by serial dilution. After overnight culture, we purified the plasmid library (pc3-MPRA-ΔEGFP) using a Maxi Plasmid Purification Kit (ThermoFisher, A31231). We digested the expanded library pc3-MPRA-ΔGFP with BamHI restriction enzyme, and inserted an amplicon containing minimal promoter with TATA box (same as Addgene plasmid 109036) and EGFP. In the reporter library pc3-MPRA-EGFP, designed enhancers or reference enhancers were upstream of the minimal promoter and the barcodes were located in the 3*^′^* UTR of EGFP. All enhancer sequences tested in Figure 3 are in the same library. We ex-panded the reporter library with Escherichia coli (XL10-Gold, Agilent Technologies) and purified it using a Maxi Plasmid Purification Kit (ThermoFisher, A31231). Both pc3-MPRA-ΔEGFP and pc3-MPRA-EGFP libraries were sequenced with 2 *×* 100 bp chemistry on Illumina MiSeq i100 plus.

To construct the enhancer AAV, we obtained a plasmid from Addgene (224004), which contained an enhancer (AiE0680m), a minimal beta-globin promoter and a SYFP2 reporter. We replaced the enhancer AiE0680m with designed or reference enhancer sequences to keep the expression cassette the same across compared enhancers and purified plasmids with a Maxi Plasmid Purification Kit (ThermoFisher, A31231).

### MPRA library construction

For the adhesive cells (HepG2, SK-N-SH), we plated them in 6-well plates (ThermoFisher, 150628) one day before transfection. We transfected cells with Lipofectamine 3000 (ThermoFisher, L3000015) following the standard protocol and we added 2 *µ*g of library plasmid each well (1 million cells). For the suspended cells (K562), we used Nucleofector 2b (Lonza) following the default protocol for K562 and we added 2 *µ*g of library plasmid per 1 million cells.

After expression for 48 hours, we extracted all RNAs from cells using an RNA MiniPrep kit (Zymo Research, R2070). Then we removed gDNA using ezDNase, and synthesized cDNA using an EGFP-specific primer with a SuperScrip IV First-Strand Synthesis kit (ThermoFisher, 18091150). We then pre-amplified the library by PCR using cycle numbers determined by qPCR. Finally, we added sequencing adaptors to the library and sequenced the library with 2 *×* 100 bp chemistry on Illumina MiSeq i100 plus.

### MPRA data analysis

DNA-library and RNA sequencing read-abundance tables were processed using custom Python scripts. Twelve-nucleotide barcodes were extracted downstream of the shared flanking sequence. Extracted barcodes were assigned to the closest designed barcode by Hamming distance, allowing at most two nucleotide substitutions. Unassignable reads were excluded from the abundance calculation. Read counts were summed for each designed enhancer–barcode pair. For each cell type (HepG2, K562, and SK-N-SH), RNA measurements from two independent transfections on two different days were retained as separate biological replicates and analyzed against one shared DNA-library reference. Elements with at least 10 reads were retained. Counts were analyzed using PyDESeq2 (Muzellec et al., 2023) with positive-count size-factor normalization and a negative-binomial generalized linear model containing sample type (RNA versus DNA) as the predictor. Enhancer activity was defined as the estimated log2 fold change of RNA relative to DNA.

### Mice

For *in vivo* experiments, wild-type C57BL/6 mice were used with procedures approved by the Stanford Institutional Animal Use and Care Committee. Mice were housed at 45%–65% humidity and 20–24*^◦^*C on a 12-hour light/dark cycle. Male and female mice were used interchangeably for the experiment. The mice were 5–6 weeks-old at the time of retro orbital injection. For each mouse, we injected 7 *×* 10^11^ vg in 110 *µ*L ice-cold PBS. We waited 3 weeks for expression before proceeding to downstream experiments.

### Histology

Once anesthetized with isoflurane, mice were perfused transcardially with ice-cold PBS and fixed with 4% paraformaldehyde (PFA) in PBS. Mouse brains were dissected and immersed in 4% PFA/PBS at 4 *^◦^*C overnight. Fixed brains were then placed in 30% sucrose in PBS for two days until it sank at 4 *^◦^*C. Brains were frozen and sliced into 50-*µ*m thick coronal sections using a sliding microtome (Leica). Brain slices were collected and mounted on slide glasses in DAPI mounting medium (VectaShield H-1500, Vector Laboratories). The slides were imaged using a fluorescent microscope Axiovert 200M (Zeiss) equipped with a 10*×* objective. The excitation light was filtered by a 514/10 nm filter and the fluorescence was collected with a 545/40 nm filter. Images were acquired with *µ*Manager 2.0 software and stitched using python scripts.

### 10x single cell sequencing

Mice were anesthetized with isoflurane and perfused transcardially with ice-cold PBS and fixed with 4% paraformaldehyde (PFA) in PBS. Once dissected, mouse brains were kept on ice and immersed in 4% PFA/PBS at 4 *^◦^*C overnight. We washed mouse brains in ice-cold PBS twice, and dissociated them with liberase (Sigma-Aldrich, 5401151001). The dissociated brains were filtered through 70 *µ*m filters to get single cells and stained by DAPI. We then sorted GFP+ cells using FACSAria Fusion, and collected 3 *×* 10^5^–1 *×* 10^6^ cells for each brain. Sorting solution and collecting solution contains RNase inhibitor (New England Biolabs, M0314L) to minimize RNA loss.

We followed the protocol for GEM-X Flex v2 (10x Genomics, CG000834) for probe hybridization, barcode oligo hybridization, washing, GEM generation and barcoding, GEM recovery and pre-amplification, and library construction. For probe hybridization, we spiked in six-pairs of customized probes into the mouse WTA probes following the Flex v2 guideline to capture SYFP2 reporter gene. For each mouse brain, we hybridized 250,000 sorted cells, and generated GEM with a subset of hybridized cells to get an estimated recovery of 50,000 cells. SYFP2 libraries and mouse genome libraries were constructed separately following the Flex v2 guideline. We mixed 10% SYFP2 library and 90% mouse genome library and conducted sequencing on Illumina NovaSeq X with 54, 10, 10, 50 reading configuration.

### Single-cell analysis and cell-type assignment

Sequenced libraries were processed with Cell Ranger 10.0.0 multi against the GRCm39-2024-A reference (GENCODE vM33) and the mouse WTA probe set v2.0.0. The SYFP2 reporter library was directly quantified from the FASTQs, counting a molecule once per unique (cell barcode, probe identity, UMI) tuple. SYFP2 counts were joined to Cell Ranger cell calls on (sample, gel-bead barcode) within each GEM well. Cell types were assigned with the Allen Institute MapMyCells cell_type_mapped against the Whole Mouse Brain 10x taxonomy CCN20230722(34 classes and 338 subclasses) with raw-count normalization, hierarchical assignment, 100 bootstrap iterations at a bootstrap factor of 0.9, a chunk size of 5,000 cells, and five runners-up retained. Assignments were confident at subclass level with median bootstrap probability of 1.000. Endothelium and the olfactory bulb were removed before analyzing the cell type distribution. In Figure 4G and S13, we gated SYFP2 positive cells based on UMI count (>100 UMIs for 6 pairs of SYFP2 probe total) and probe appearance (>5 pairs out of 6). We have also shown the SYFP2 UMI distribution and how the threshold affects the SYFP2 positive cell composition in Figure S14.

## Data and Code Availability Statement

All source data will be made available from the corresponding authors upon request. Scripts for genomic ICL with Evo 2 and post-training are in the GitHub repo (https://github.com/evo-design/ICL_regulatory.git).

## Acknowledgements

We thank the Stanford Gene Vector and Virus Core for viral vector packaging, and Arc Institute for flow cytometry and NGS sequencing services. We thank Brian Plosky for helpful discussions and assistance with manuscript preparation. Y.A.H. acknowledges Pengli Wang (Stanford University) for sharing the AAV.CAP.B10 capsid plasmid, Sungmoo Lee, Daesun Song, Yiting Wang and John Kochalka (Stanford University) for training in mouse procedures. We thank Michael Lin and Jun Ding for allowing us to use the microscope and the microtome. We thank all members of the Laboratory of Evolutionary Design for helpful discussions.

B.L.H. acknowledges funding support for this project from the Silicon Valley Community Foundation, as well as broader research support from the Gates Foundation, the NIH Director’s New Innovator Award, the Advanced Research Projects Agency for Health (ARPA-H), Arc Institute, Schmidt Sciences AI2050, Stanford Center for Digital Health, and Stanford Human-Centered Artificial Intelligence (HAI) Hoffman-Yee Research Grants. M.L. acknowledges funding support from the Stanford HAI Data Science Scholars program.

G.B. acknowledges funding support from the National Science Foundation Graduate Research Fellowship Program. T.L. acknowledges funding support from the Stanford Knight-Hennessy Scholars program. D.C. acknowledges funding support from the Stanford Genomics Training program.

## Author Contributions

Y.A.H. and B.L.H. conceived the project. B.L.H. supervised the project and contributed to the code for genomic ICL. Y.A.H. developed the code for genomic ICL, Evo 2 post-training, and performed the *in vitro* and *in vivo* experiments. M.L. performed Categorical Jacobian and motif density analysis. D.N. contributed to the code for genomic ICL, and developed code for DPO post training. G.B. contributed to the code for genomic ICL, and developed code for CFG in Evo 2. D.C. performed SAE analysis. R.P. contributed to Evo 2 post-training. T.L. and C.R. contributed to *in vivo* experiments under the supervision of W.E.A.

## Supplementary Figures

**Fig. S1.**
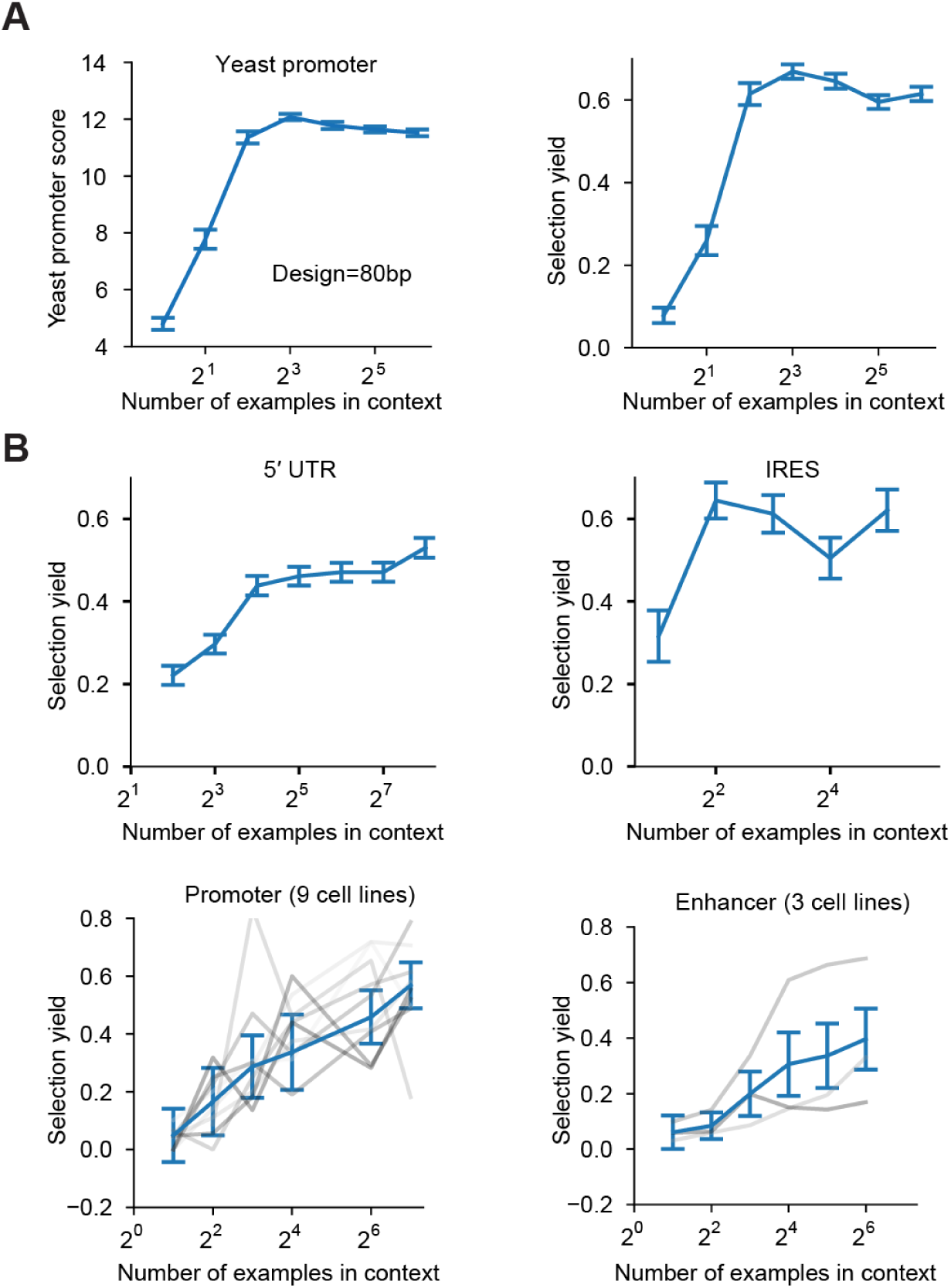
*|* **Selection Yields of Evo 2 in-context learning. (A)** Predicted activity (left) and selection yield (right) of sequences generated by Evo 2 through in-context learning for design of yeast promoters. **(B)** Selection yields of sequences generated by Evo 2 through in-context learning for four tasks: human 5*^′^* UTR design, human IRES design, human promoter design, and human enhancer design. Error bars indicate the standard error of the mean (SEM; n = 128 sequences per condition). For human promoters, gray lines represent the nine cell lines, ordered from light to dark: AGS, HAP1, HCT116, HEK293, HepG2, K562, LNCaP, MCF7, and U2OS. For human enhancers, gray lines represent the three cell lines, ordered from light to dark: HepG2, K562, and SK-N-SH.

**Fig. S2.**
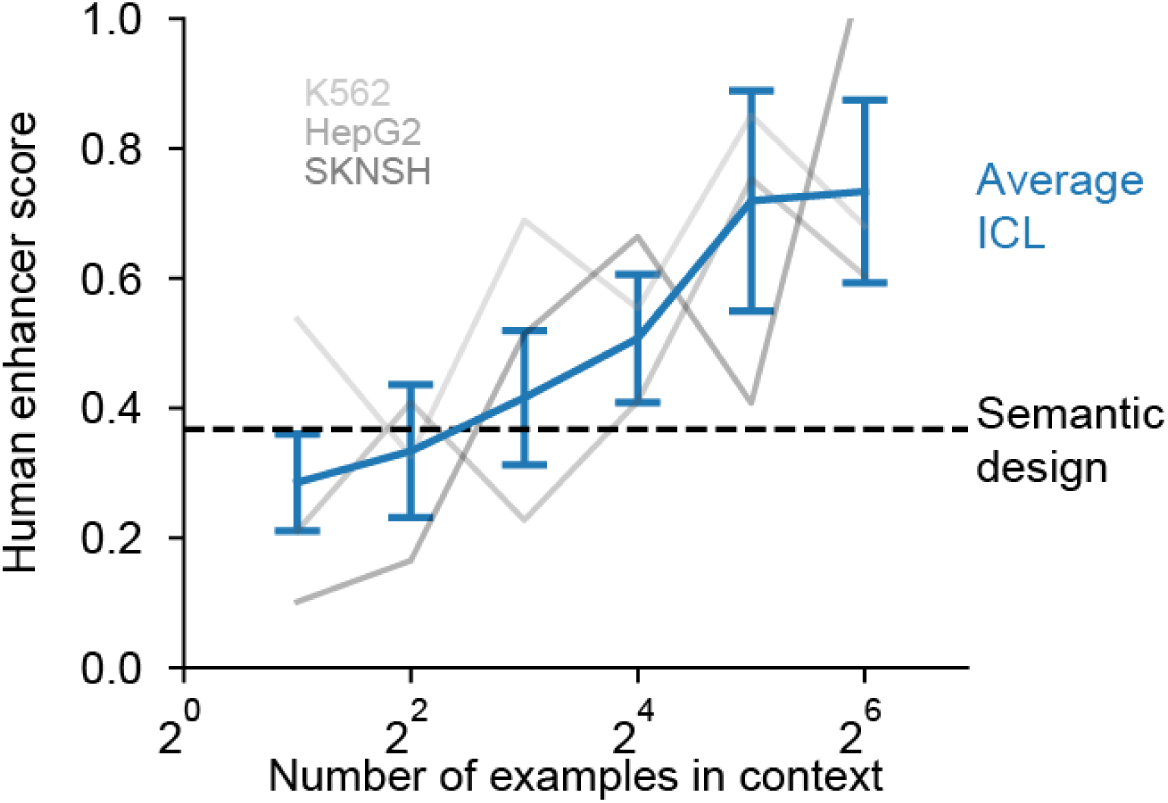
*|* **Genomic ICL and semantic design of distal regulatory elements in Evo 2.** Evo 2 can learn sequence design with better performance than semantic design for distal regulatory sequences. For semantic design, we prompted Evo 2 by the genomic context upstream of an enhancer. Here the error bar indicates SEM (n = 384 sequences per condition). See also Figure S12.

**Fig. S3.**
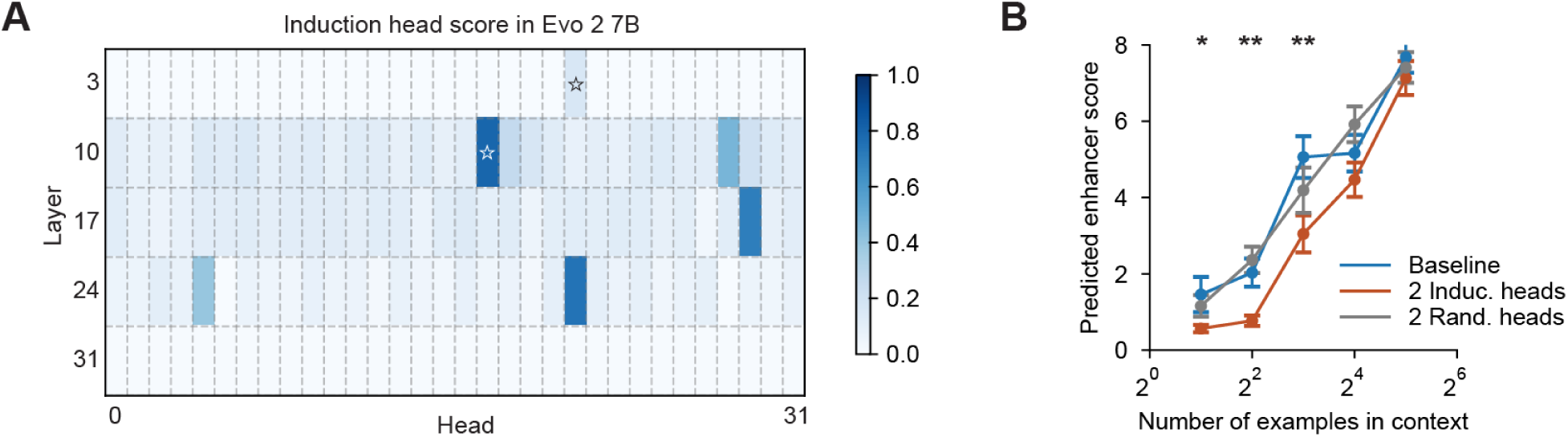
*|* **Identification and ablation of induction heads in Evo 2 7B. (A)** Identification of induction heads in all 5 attention layers in Evo 2 7B. Stars indicated two induction heads ablated in Evo 2 7B. **(B)** Ablating the two induction heads impacted ICL in Evo 2 7B when compared to two randomly selected heads in the same layers. Error bars indicate SEM (n = 32 ICL prompts per condition from). The *p*-values for two-sided paired *t* -test are 0.0499 (2-shot), 0.0046 (4-shot), 0.0097 (8-shot) corrected by the Benjamini-Hochberg procedure.

**Fig. S4.**
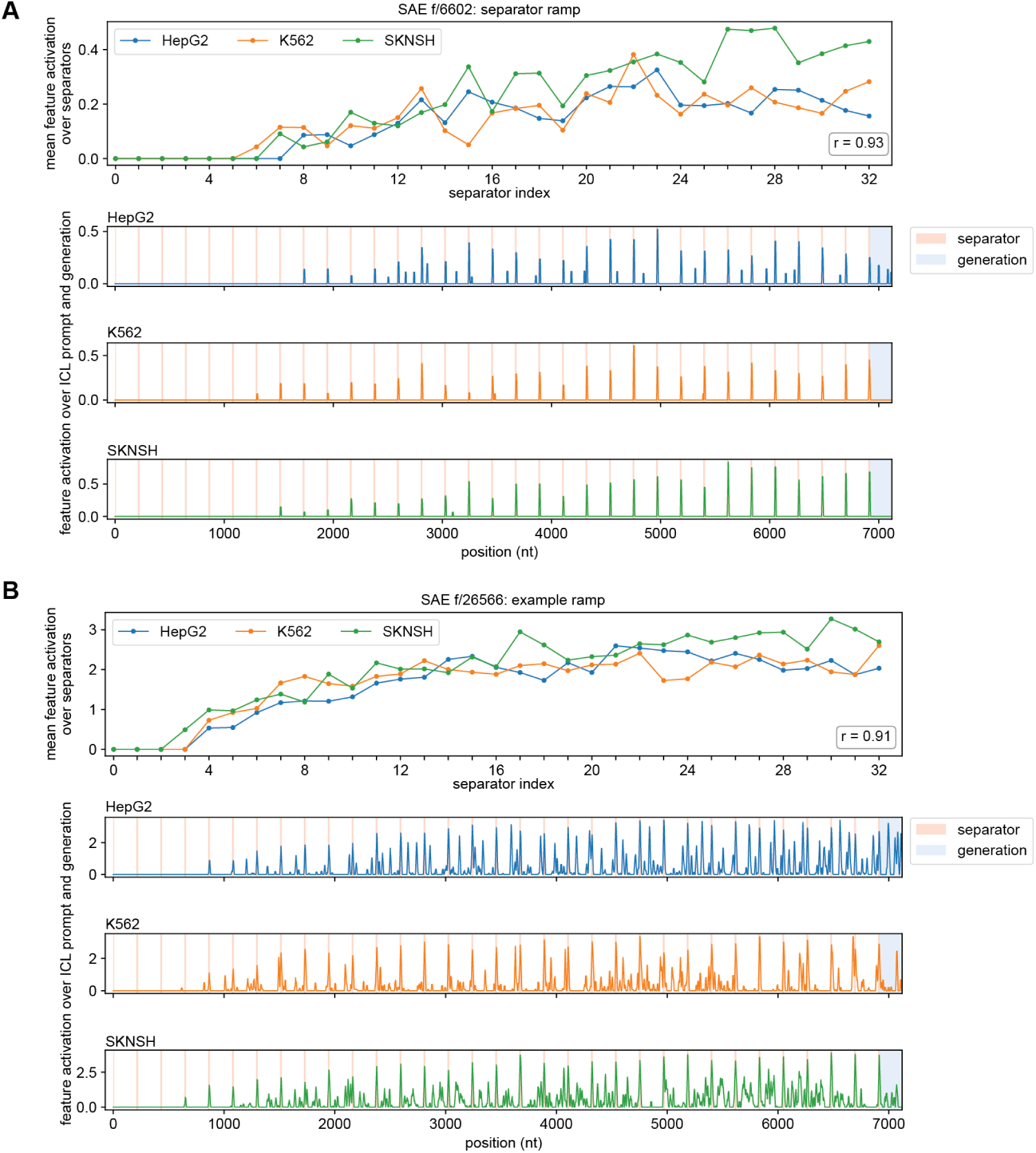
*|* **Evo 2 activation patterns track in-context learning. (A)** Concatenated example enhancers and the corresponding generated enhancer were fed into Evo 2 to extract features from a trained sparse autoencoder (SAE). SAE feature f/6602 is activated at separator sequences and increases as Evo 2 reads more enhancer examples. f/6602 shows this pattern across three cell lines. Top: integrated SAE feature activation within each separator window across successive separators. Bottom three panels: SAE feature activation across the concatenated sequences for enhancers from three cell types. **(B)** SAE feature f/26566 is activated at both separator sequences and enhancer sequences, and increases as Evo 2 reads more enhancer examples. Panel layout is the same as in **(A)**.

**Fig. S5.**
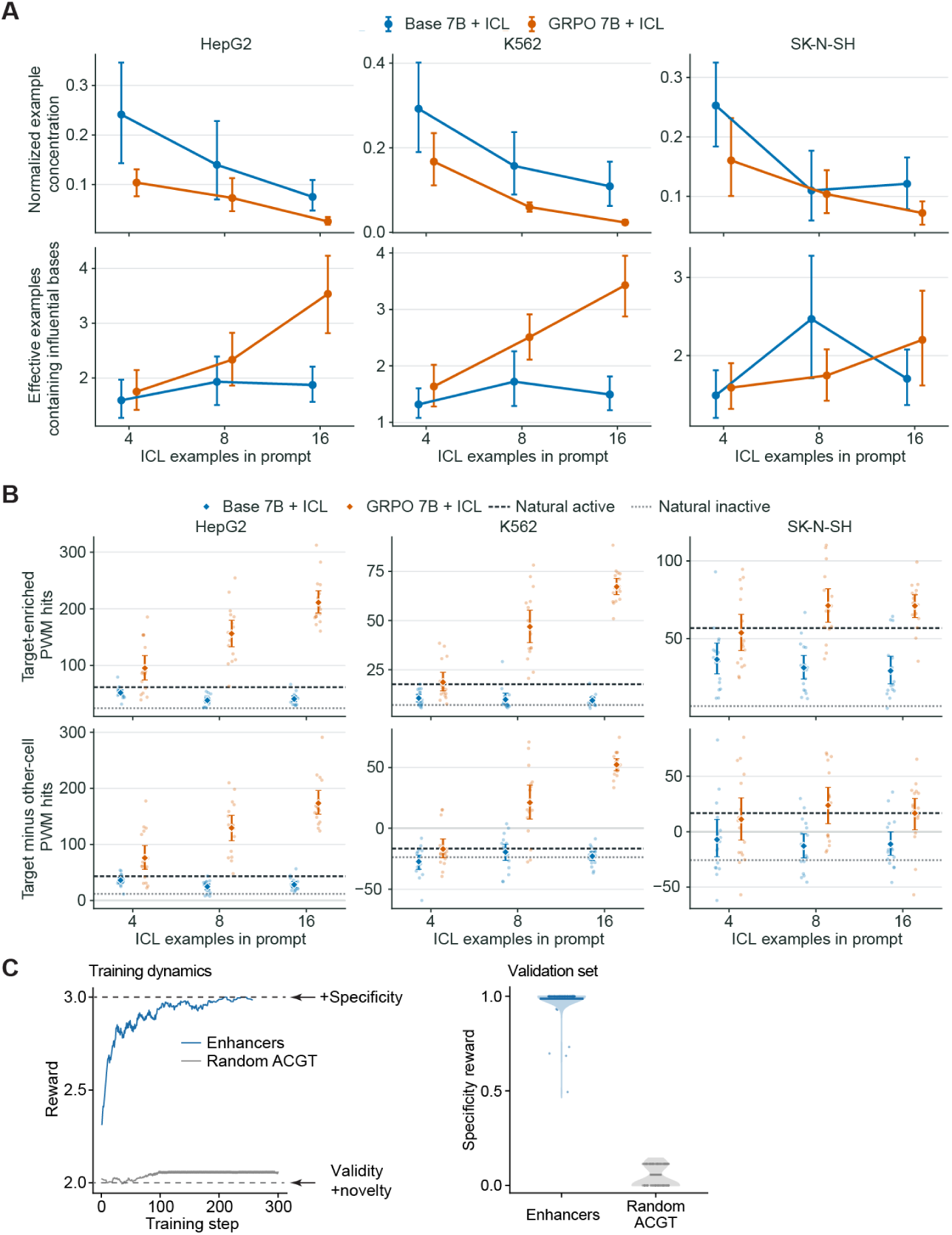
*|* **Mechanistic analysis of post-trained Evo 2. (A)** Distributed influence of in-context examples. Top panels: Concentration of influence from individual example sequences on generated sequences, calculated from raw categorical Jacobian contacts. Here a concentration of 1.0 indicates a single example dominates the generation, 0.0 indicates equal contribution from all examples. Points are means across 16 prompt-generation pairs per condition, and error bars are 95% bootstrap intervals obtained by resampling prompts 4,000 times. Bottom panels: Effective number of examples represented among the 40 most influential bases among categorical Jacobian contacts. Here an effective support of 1 indicates all 40 most influential bases come from one example sequence, larger values indicate these bases are distributed across more examples. **(B)** Motif density across number of examples in context. Top panels: Target cell type-enriched PWM hits for each generated sequence before and after GRPO. Dots indicate the mean of eight generations from one prompt; diamonds are means across all 16 prompts. Error bars are 95% bootstrap intervals obtained by resampling prompts 4,000 times. Means of natural sequences predicted to be active and inactive in each target cell type are respectively shown as dashed and dotted lines. Bottom panels: On-target minus off-target PWM hits for each generated sequence before and after GRPO. Counts are not adjusted for the number of enriched PWMs in each cell type. **(C)** Effects of ICL in post training. Left: Dynamics of post-training reward with enhancer ICL or random sequence ICL. Note that valid and novel ACGT generations with proper GC content can have a reward of 2.0. The additional reward is for specificity. Right: Specificity reward of sequences generated using ICL with the validation set.

**Fig. S6.**
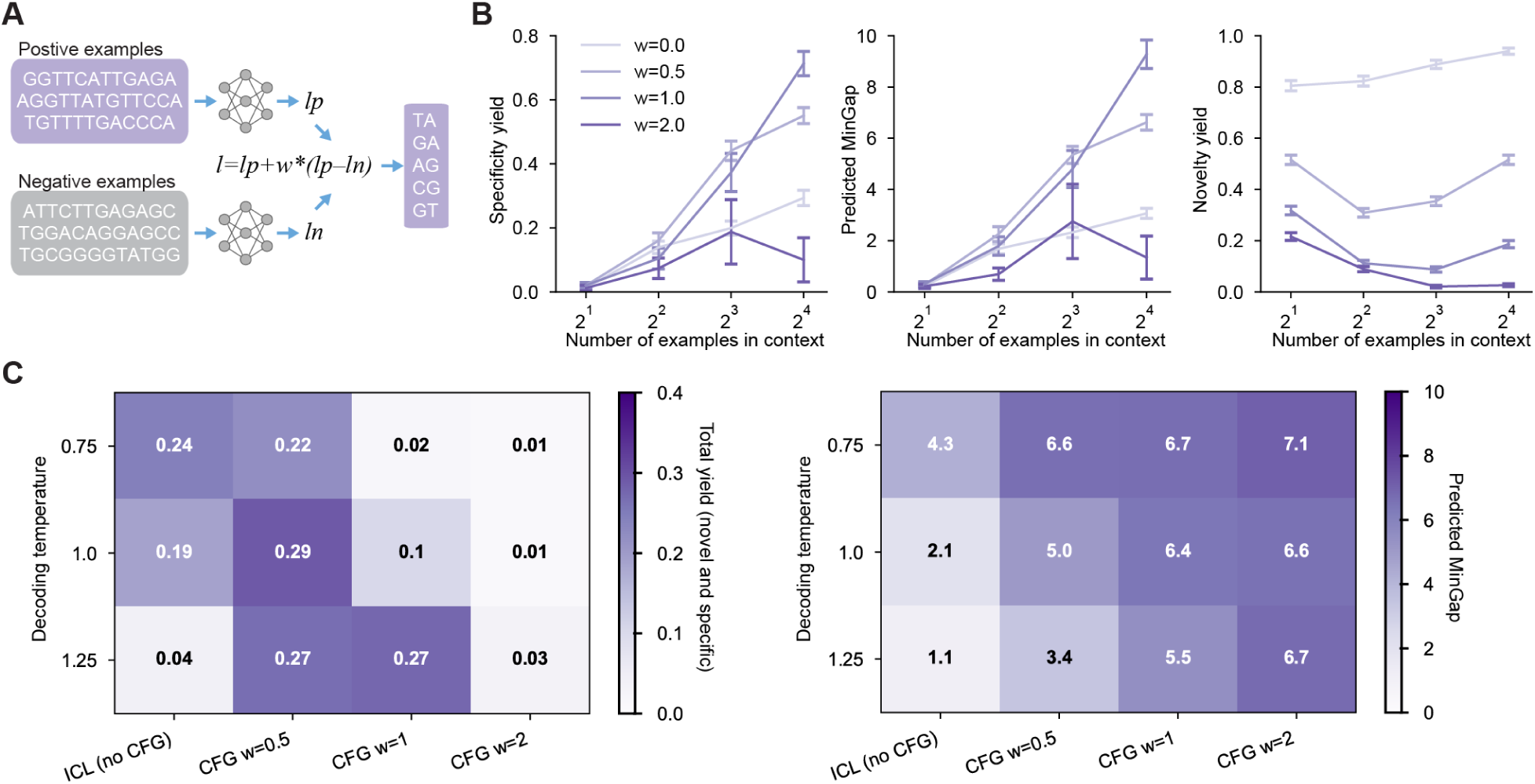
*|* **ICL with classifier free guidance. (A)** Schematic of classifier-free guidance of sequence design with positive and negative examples. **(B)** Specificity yield (left), predicted MinGap (middle) and novelty yield (right) as a function of steering strength w and number of in-context examples. Steering of intermediate strength increases specificity at the cost of sequence novelty. **(C)** Total yield (left) and predicted MinGap (right) as a function of steering strength and decoding temperature. High steering strength requires high decoding-temperature to achieve high overall yield, although it achieved higher specificity at lower temperature.

**Fig. S7.**
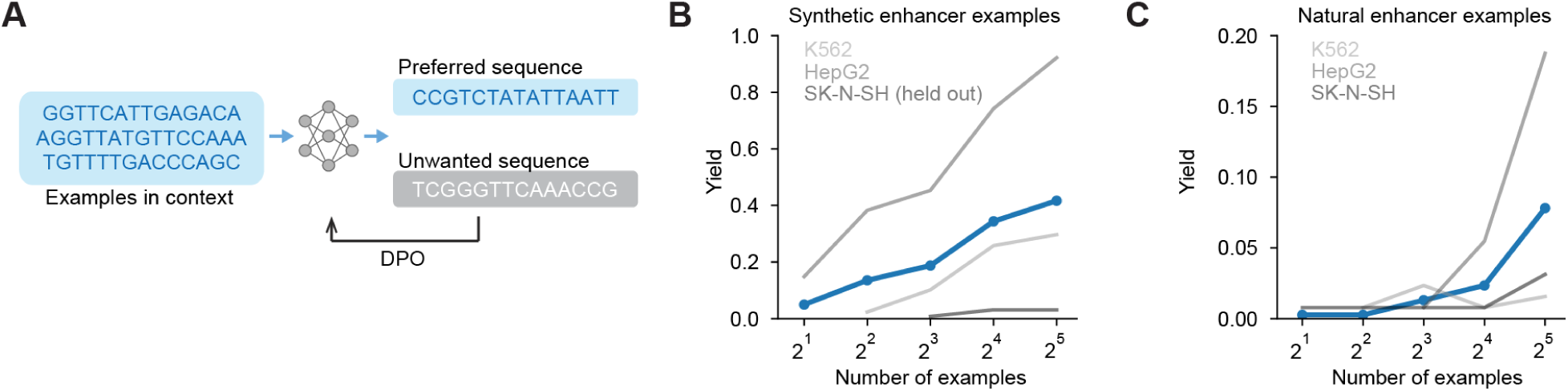
*|* **Direct preference optimization in Evo 2 with ICL. (A)** Schematic of direct preference optimization in Evo 2. **(B)** Fine-tuned Evo 2 shows higher performance with more synthetic enhancer examples in-context. **(C)** Fine-tuned Evo 2 shows higher performance with more natural enhancer examples in-context.

**Fig. S8.**
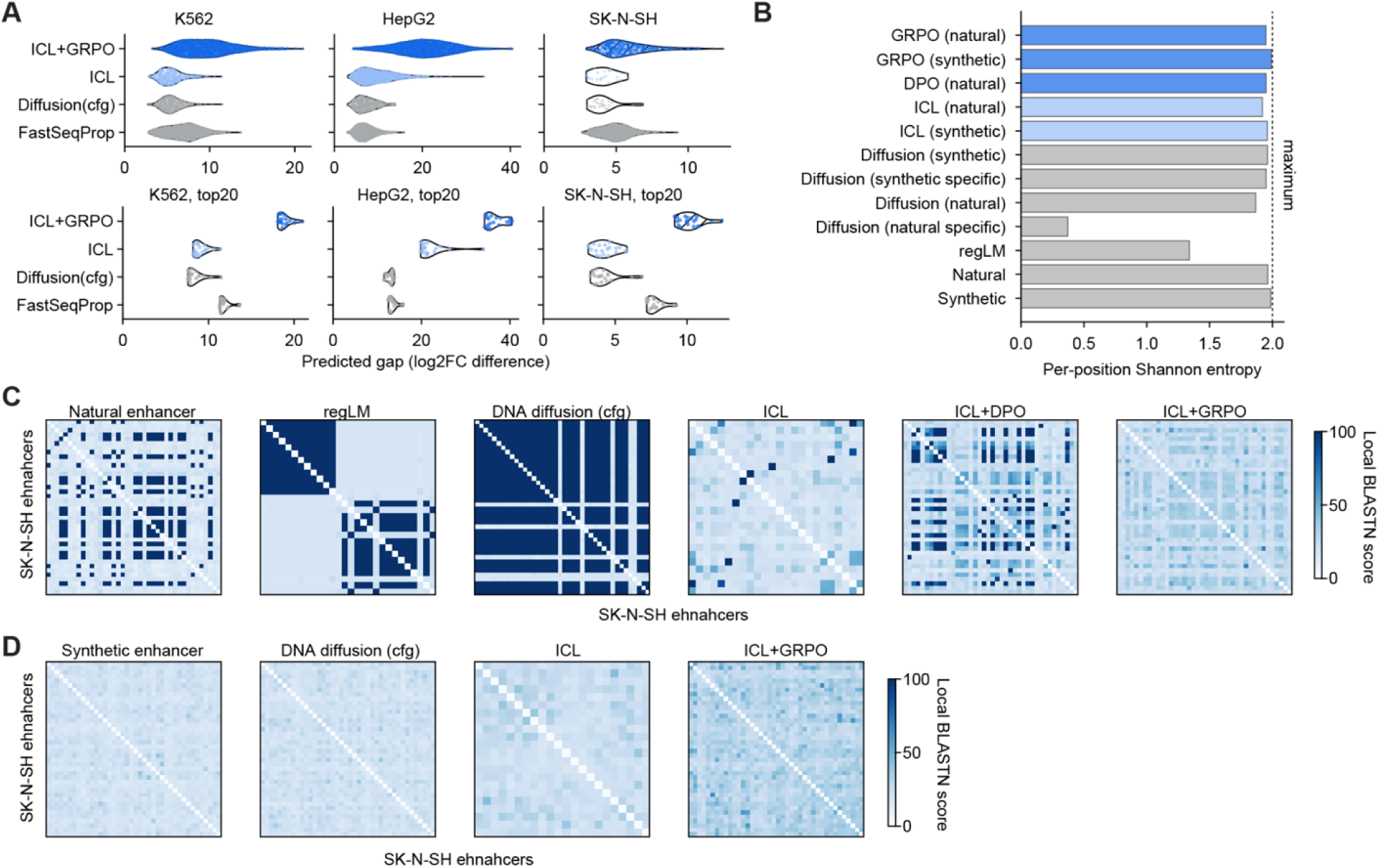
*|* **Additional benchmarking of Evo 2 in-context learning against alternative methods. (A)** Comparing Evo 2-based methods against alternative methods for predicted specificity of designing enhancer sequences, measured by MinGap. Evo 2-based methods include ICL design and ICL + GRPO, reinforced with virtual rewards. Here Evo 2-based methods use synthetic enhancers as in-context examples. Other methods include DNA diffusion with classifier-free guidance (cfg) trained on synthetic enhancer sequences; and synthetic enhancer sequences designed by a gradient method FastSeqProp. Top panels: specificity gap of filtered sequences from a fixed number of generated sequences. Bottom panels: specificity gap of the top 20 sequences. Hatching indicates SK-N-SH is held out from Evo 2 reinforcement post-training. **(B)** Comparing Evo 2-based methods with other methods for sequence diversity using per-position Shannon entropy. 2.0 is the theoretical maximum. In Figure 2 we characterized the DNA-diffusion models with CFG trained on cell-type specific synthetic enhancers, and all natural enhancers, as the model degenerated when trained only on cell-type specific natural enhancers. **(C)** Pairwise local BLASTn score in filtered SK-N-SH enhancers across 5 generative methods based on natural enhancer dataset. A BLASTn score of 60 corresponds to e-value *∼*10*^−^*^5^ and lower score means higher diversity. **(D)** Pairwise local BLASTn score in filtered SK-N-SH enhancers across 3 generative methods based on synthetic enhancer dataset.

**Fig. S9.**
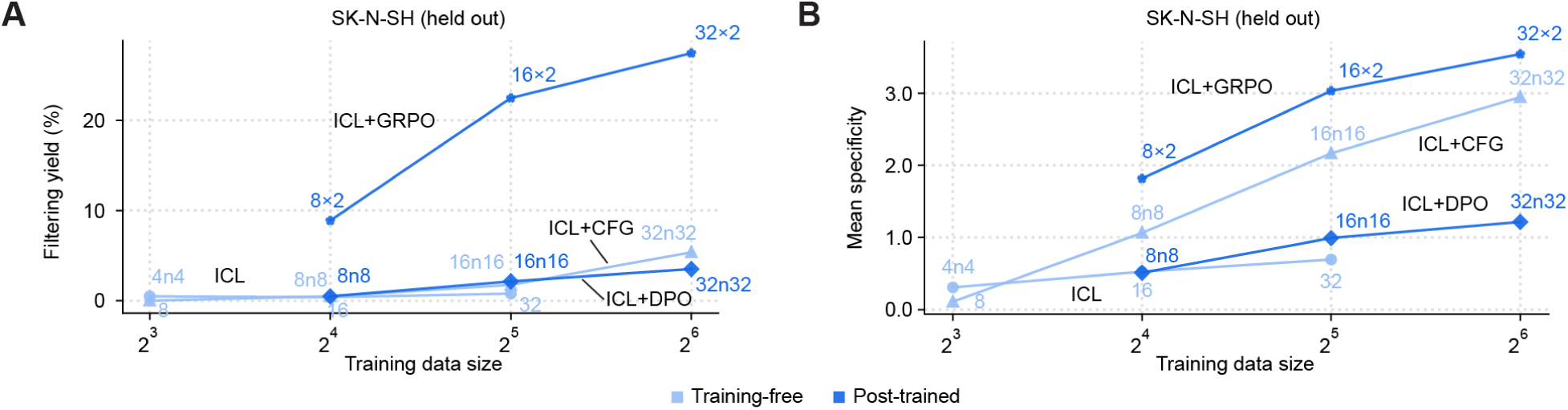
*|* **Data-efficiency on the cell type held out from post-training. (A)** Additional data for the data efficiency experiment in Figure 2F, G on a third cell type, SK-N-SH, which was held out from the Evo 2 post training, i.e. ICL+GRPO and ICL+DPO. For the training-free methods (light blue), i.e. ICL and ICL+CFG, the data size indicates the number of in-context examples of SK-N-SH enhancers. For the post-training methods (blue), the data size indicates the number of sequences used in training on the other two cell types (HepG2 and K562). Yield is defined as the proportion of generated sequences that are novel, valid, and specific to SK-N-SH. **(B)** Mean cell-type specificity of novel sequences generated by the computational frameworks evaluated in (A), across different training dataset sizes.

**Fig. S10.**
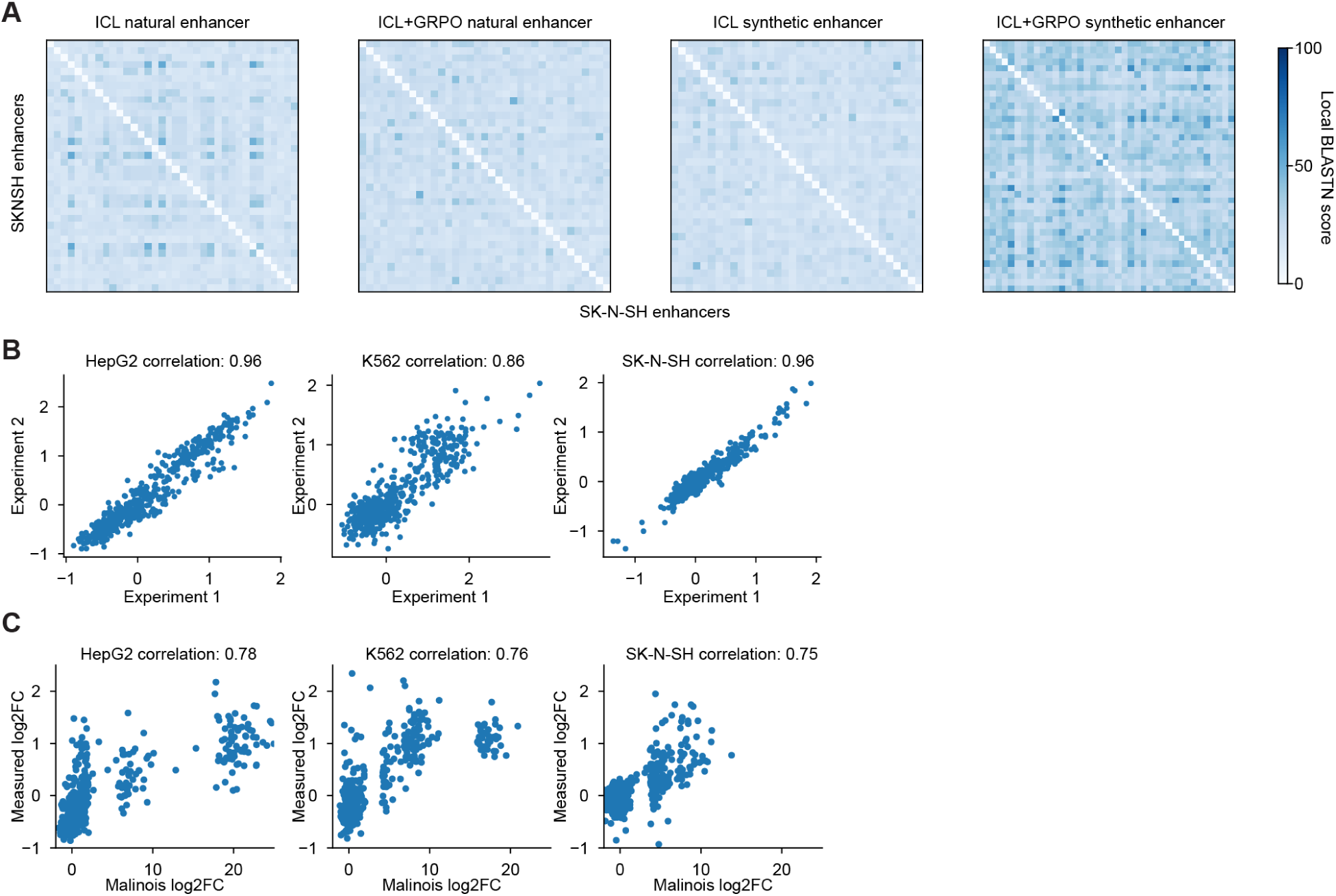
*| **In vitro*** **measurement of selected cell-type specific enhancers. (A)** Pairwise local BLASTn score in filtered SK-N-SH enhancers across four groups. A BLASTn score of 60 corresponds to e-value *∼* 10*^−^*^5^ and lower score means higher diversity. **(B)** Two independent experimental repeats show high correlations across three cell lines. **(C)** Measured log2 fold change has high correlations with predicted log2 fold change across three cell lines.

**Fig. S11.**
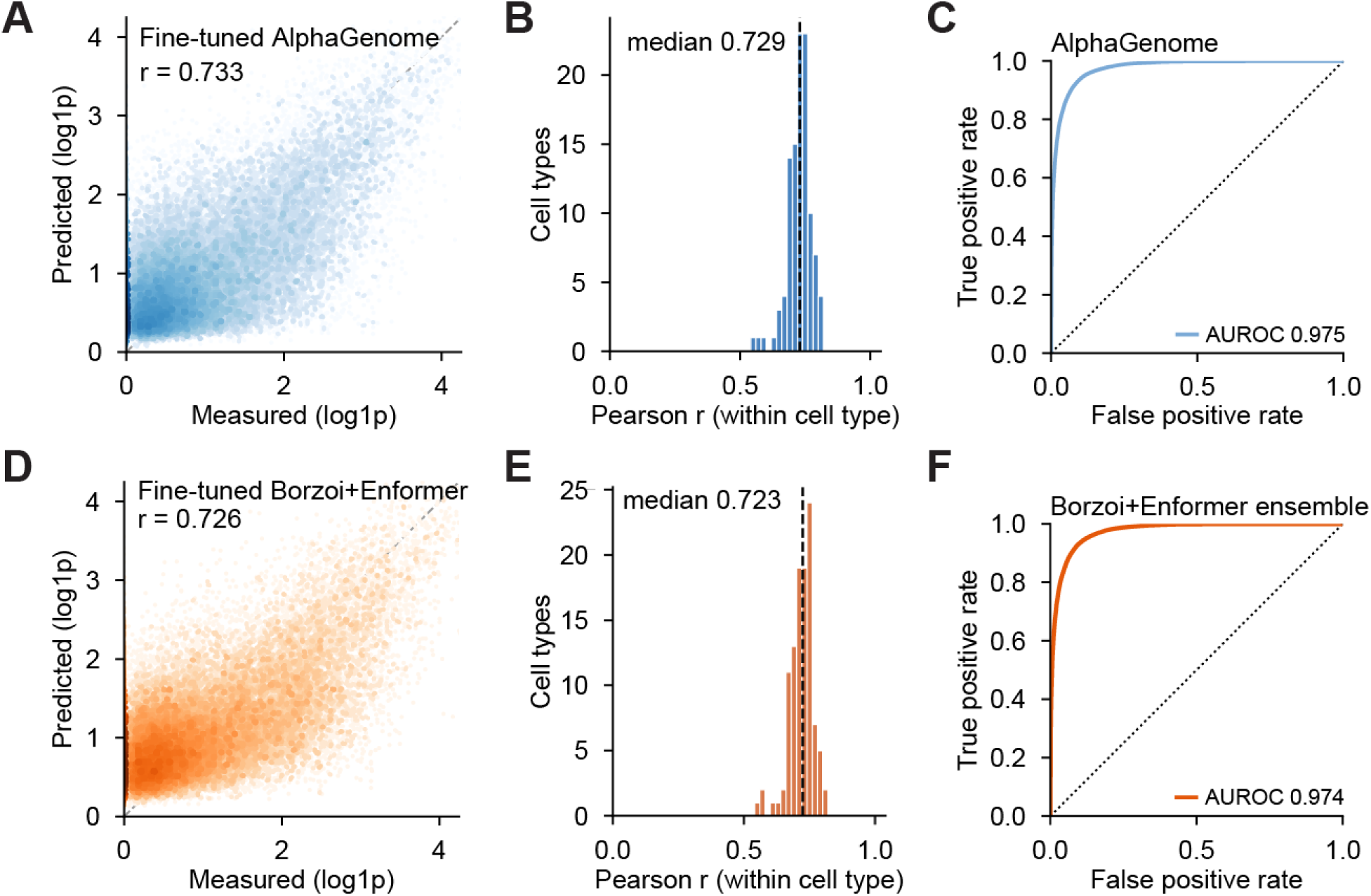
*|* **Fine-tune AlphaGenome and Borzoi ensemble to classify and predict human brain cCRE activity across 107 cell types. (A)** Scatter plot of log(1+x)-transformed measured ATAC signal and predicted ATAC signal by fine-tuned AlphaGenome on the test split, across 107 cell types. **(B)** Pearson correlation coefficients of measured and AlphaGenome-predicted ATAC signals across 107 cell types. **(C)** ROC curve of fine-tuned AlphaGenome CRE classifier on the test split. **(D)-(F)** Same as (A)-(C), where the model is the fine-tuned Borzoi+Enformer ensemble.

**Fig. S12.**
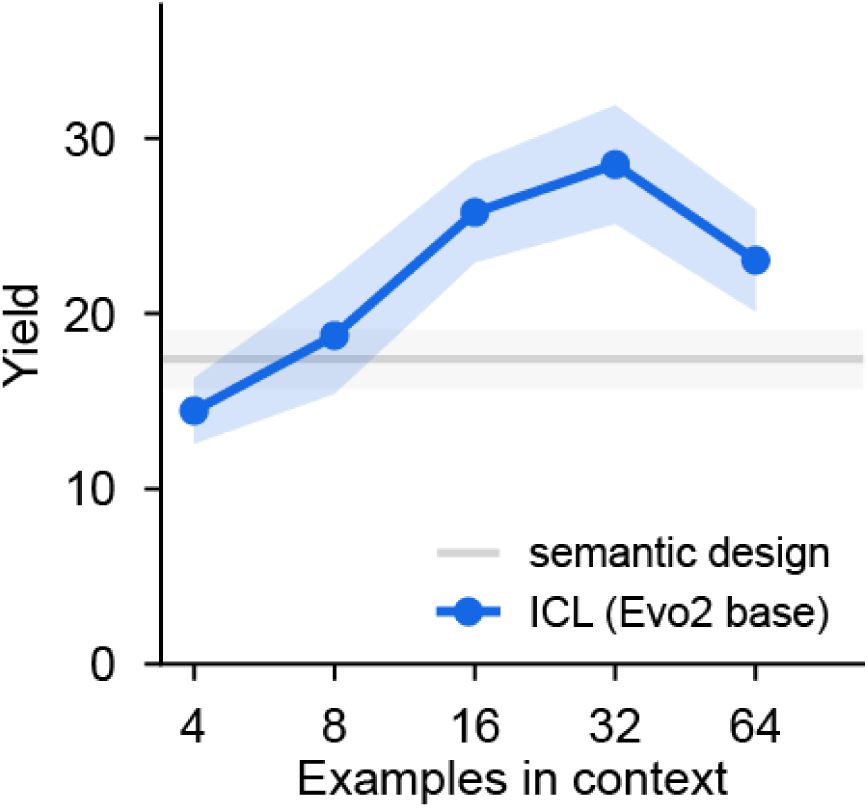
*|* **Evo 2 generates cCREs for human neurons.** Base Evo 2 model generated sequences by ICL with higher yield than by semantic design. Here, the yield is defined as the proportion of generated sequences that are novel and with classifier score > 0.5. For semantic design, we prompted Evo 2 by the genomic context upstream of a cCRE. Shadow indicates standard error of the mean across four selected cell types each with eight different prompts (n=32 prompts each condition).

**Fig. S13.**
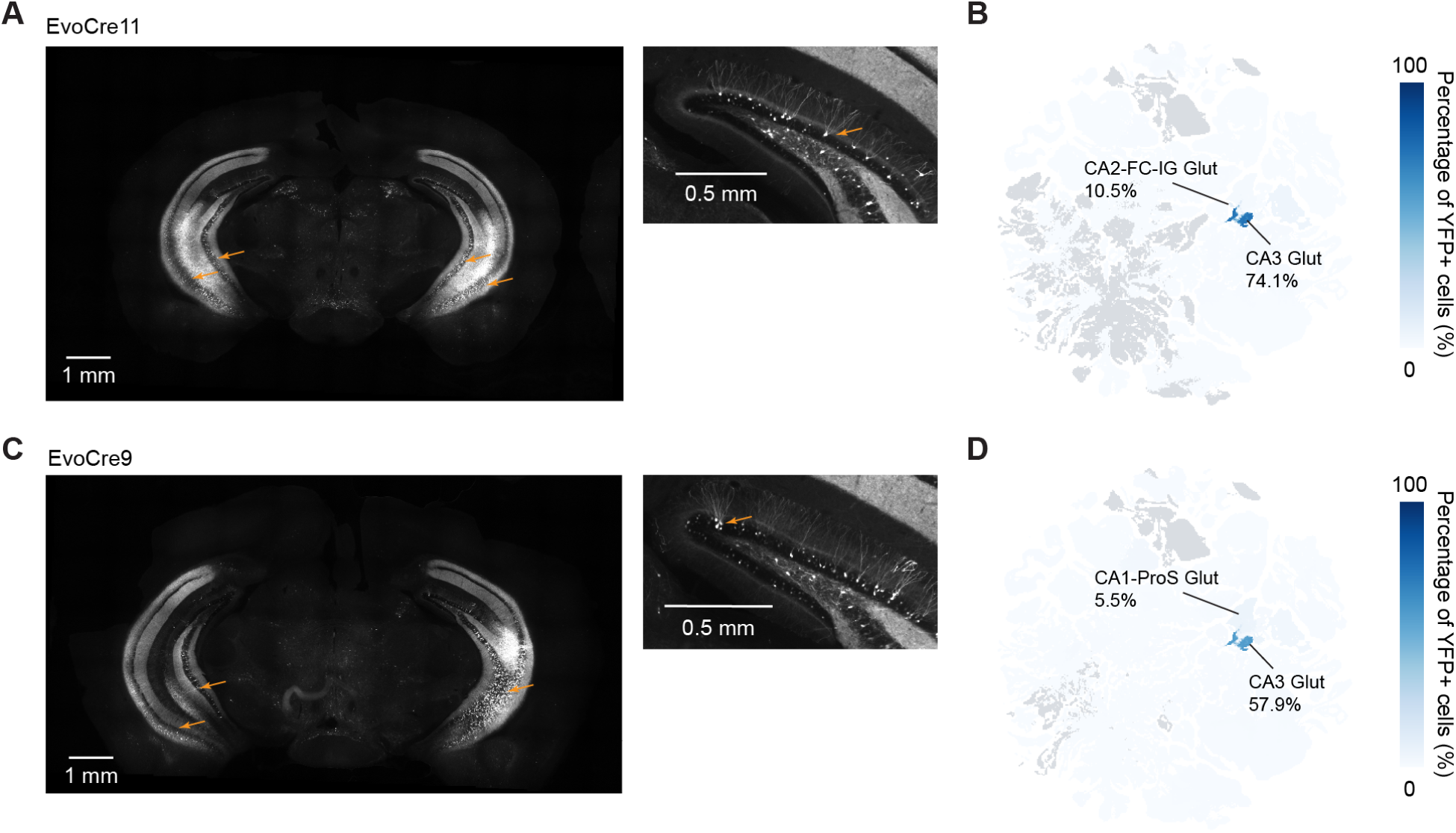
*|* **Two designed enhancers targeting CA3 excitatory neurons.** (A) Left: Global expression pattern of EvoCre11-targeted cells in a mouse brain. Orange arrows indicate CA3 excitatory neurons. Right: Expression pattern of EvoCre11-targeted cells in mouse hippocampus. Orange arrow indicates a neuron with morphology of a Vip+ interneuron in the hippocampus. (B) Distribution YFP+ positive cells across cell types for EvoCre11. Top two cell types and their proportions are annotated. Gray indicates cell types below detection threshold in the data. (C)-(D) Similar to (A) and (B) for quantifying designed enhancer EvoCre9.

**Fig. S14.**
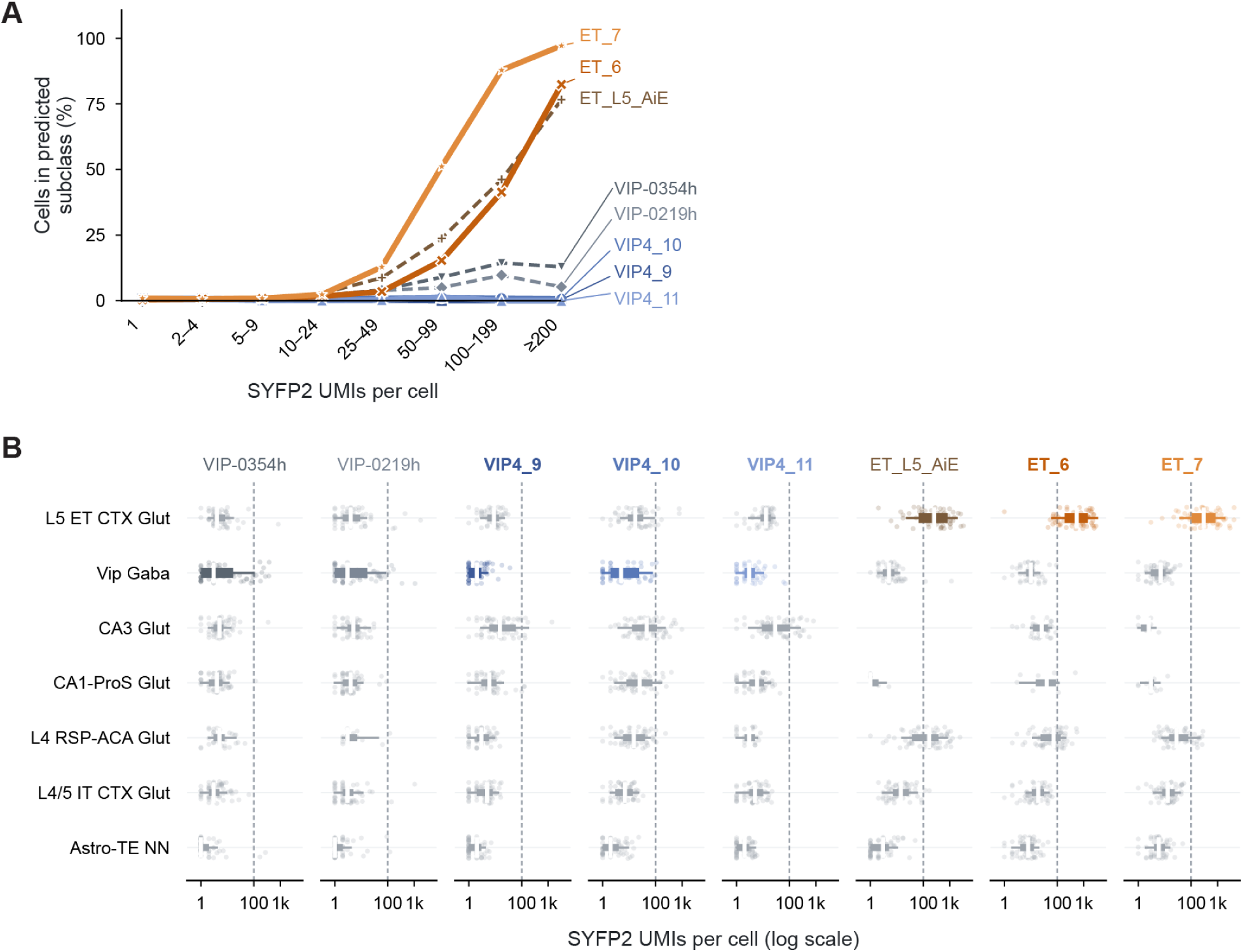
*|* **SYFP2 UMI distribution across cell types by brain-wide single-cell sequencing.** (A) Percentage of on-target cell type in YFP+ cells in different SYFP2 UMI bins. (B) Distribution of SYFP2 UMI in selected cell types across designed and existing enhancers. Dashed line at 100 indicates the threshold used in Figure 4 and S13 for quantifying the specificity.

